# CROssBARv2: A Unified Computational Framework for Heterogeneous Biomedical Data Representation and LLM-Driven Exploration

**DOI:** 10.64898/2026.04.12.718028

**Authors:** Bünyamin Şen, Erva Ulusoy, Melih Darcan, Mert Ergün, Sebastian Lobentanzer, Ahmet S. Rifaioglu, Dénes Türei, Julio Saez-Rodriguez, Tunca Doğan

**Affiliations:** Biological Data Science Lab, Dept. of Computer Engineering, Hacettepe University, Ankara, Turkey; Dept. of Bioinformatics, Graduate School of Health Sciences, Hacettepe University, Ankara, Turkey; Dept. of Health Informatics, Institute of Informatics, Hacettepe University, Ankara, Turkey; Heidelberg University, Faculty of Medicine, and Heidelberg University Hospital, Institute for Computational Biomedicine, Heidelberg, Germany; Institute of Computational Biology, Computational Health Center, Helmholtz Center, Munich, Germany; European Molecular Biology Laboratory, European Bioinformatics Institute (EMBL-EBI), Hinxton, Cambridgeshire, UK

## Abstract

Biomedical discovery is hindered by fragmented, modality-specific repositories and uneven metadata, limiting integrative analysis, accessibility, and reproducibility. To address these challenges, we present CROssBARv2, a provenance-rich biomedical data-and-knowledge integration platform that unifies heterogeneous sources into a maintainable, scalable system. By consolidating diverse data types into an extensive knowledge graph enriched with standardised ontologies, rich metadata, and deep learning–based vector embeddings, CROssBARv2 alleviates the need for researchers to navigate multiple siloed databases and can facilitate downstream tasks, including predictive modelling and mechanistic reasoning, enabling applications such as drug repurposing and protein function prediction. The platform offers interactive graph exploration and embedding-based semantic search with CROssBAR-LLM, an intuitive natural language question-answering system that grounds large language model (LLM) outputs in the underlying knowledge graph to mitigate hallucinations. We assess CROssBARv2 through (i) multiple use-case analyses to test biological coherence and relational validity; (ii) knowledge-augmented biomedical question-answering benchmarks comparing CROssBAR-LLM against generalist LLMs; and (iii) a deep learning–based predictive modelling experiment for protein function prediction leveraging the heterogeneous structure of CROssBARv2. Collectively, CROssBARv2 provides a scalable, AI-ready, and user-friendly foundation that facilitates hypothesis generation, knowledge discovery, and translational research.

## Introduction

Biomedical discovery increasingly depends on integrative, data-driven approaches to elucidate disease mechanisms and accelerate therapeutic development. Yet biological knowledge is scattered across isolated resources, hindering the discovery of meaningful relationships and preventing the full exploitation of complementary information ^1^. Overcoming this fragmentation demands unified and robust frameworks that explicitly represent heterogeneous biological entities and their relationships, enabling analysis of complex interactions and uncovering insights elusive in isolated datasets.

Knowledge graphs (KGs) meet this need by modelling biological entities as nodes, and their interactions as edges. KGs integrate diverse datasets into a flexible and scalable network that preserves contextual relationships and support computational analyses such as pathfinding and link prediction. Several efforts exemplify current practice: Hetionet interlinks several biological entities to prioritise drug-repurposing candidates ^2^; SPOKE bridges molecular findings and clinical concepts, facilitating translational inference ^3^; PrimeKG integrates 20 curated sources for precision-medicine queries ^4^; Bioteque aggregates >150 datasets to create pre-computed embeddings for machine-learning tasks ^5^; the Clinical Knowledge Graph (CKG) consolidates biomedical databases, experimental results, and scientific literature for clinical proteomics and omics workflows ^6^; and CROssBARv1, the precursor to this study, integrates large-scale biomedical data from 14 databases together with machine learning-based predictions into a KG with a graphical web interface for intuitive data exploration and interpretation, focusing on drug discovery ^7^. Finally, the BioCypher framework automates FAIR-compliant ingestion, harmonisation, and export of biomedical data—complete with provenance—into graph databases such as Neo4j ^8^.

While progress has been made in unifying fragmented data within integrated platforms, significant limitations remain in functionality, maintainability, reproducibility, and accessibility. Many current KGs are designed for highly specialised tasks (e.g., DRKG and PrimeKG), which limits their flexibility and constrains their utility for broader research applications. In addition, existing KGs often lack comprehensive metadata for biological entities, neglecting valuable contextual information such as data provenance, experimental validation status, and confidence scores that could enhance their reliability, utility, and enable more informed decision-making. Without such metadata, users may conflate curated and predicted interactions, leading to potentially erroneous biological conclusions. Usability is another key issue. Many KGs are distributed in formats that require programming skills for data extraction and analysis, restricting usability for non-programmers. Reproducibility is also a significant concern. Numerous influential KGs, including DRKG and PharmKG ^9^, do not publicly share their construction code, hindering independent replication and community-driven improvements. A further limitation lies in how KGs represent entities. While several studies have applied KG embeddings ^5,10,11^, none have exploited embeddings from external pretrained models (e.g., ESM2, ProtTrans), which encode biologically meaningful features and enable retrieval of functionally similar entities even without direct links. Addressing these gaps is crucial for developing adaptable, user-friendly, and sustainable KG-based biomedical systems.

Large language models (LLMs) have emerged as powerful tools for processing and generating natural language, with broad applications in biology and biomedicine. However, generalist LLMs give users limited control over generated content and are prone to hallucinating spurious information, and many still lack direct access to specialised databases ^12,13^. To address these limitations, several strategies have been developed, including agentic web search ^14^, and Model Context Protocol (MCP) ^15^. Another approach is to integrate LLMs with domain-specific KGs. By combining the linguistic capabilities of LLMs with the rich repository of structured, interconnected, and up-to-date data in KGs, it is possible to enhance the accuracy, depth, and reliability of question-answering systems, enabling more context-aware, robust, and trustworthy applications in biomedical research ^16–18^. The extended version of this introduction, together with a detailed discussion of the relevant literature, is provided in **Supplementary Information S1**.

In this study, we present a comprehensive computational framework, CROssBARv2, designed to integrate large-scale, heterogeneous biological and biomedical data from diverse sources into a unified KG. Our reproducible pipelines systematically consolidate information from 34 data sources, encompassing 14 node and 51 edge types. We also incorporate precomputed embeddings from state-of-the-art biomedical representation models as node properties. The CROssBARv2 is deployed on a user-friendly web platform that supports fast querying and interactive visualisations via the Neo4j Browser, a GraphQL API for programmatic data retrieval, and natural language-based interaction capabilities through its integrated LLM component, CROssBAR-LLM, enabling easy exploration and interaction with the data.

Our key contributions include: (i) a provenance-rich, general-purpose KG enhanced with comprehensive metadata (provenance, evidence, confidence scores) to support reliable, filterable analyses across diverse biomedical applications; (ii) automated and maintainable data integration pipelines that ensure periodic updates and seamless incorporation of new data sources; (iii) accessible interfaces for both non-programmatic and programmatic users; (iv) an intuitive online platform that enables natural language interaction and exploration of the KG; and (v) semantic similarity search functionality using integrated node embeddings that not only enable exploration beyond topological connections, but also provide machine learning-ready feature representations for downstream models.. CROssBARv2 is designed to support a broad scientific community. Biologists and clinicians can use its visual interface and natural language–based querying tool to explore complex systems, validate drug targets, and investigate disease mechanisms without sifting through multiple databases or publications. Meanwhile, bioinformaticians and computational biologists can directly employ the KG and its embeddings for large-scale network analyses, predictive modelling, and workflow development in applications such as drug discovery, biomolecular function prediction, and disease mechanism inference. Overall, the framework aims to accelerate data-driven discovery in biomedical research.

To assess the utility and biological relevance of our KG, we performed multiple use-case analyses, evaluating selected biomedical relationships within the graph against existing literature. We also benchmarked several LLMs in terms of their potential utilisation in biomedical research and compared the performance of standalone LLMs against CROssBAR-LLM in scientific question answering. Furthermore, we evaluated the utility of the heterogeneous biomedical knowledge captured in CROssBARv2 for predictive modelling by training deep learning models for protein function prediction.

## Results

### The Overview of CROssBARv2

An overview of the CROssBARv2 workflow is provided in **Figure 1**. CROssBARv2 integrates biological data from 34 biological data sources (**Figure 1a**) into a single, unified framework that establishes biologically relevant relationships to support a wide range of analyses, using the knowledge graph data structure. The system employs configurable and reusable adapter scripts to automate the retrieval, standardisation, and integration of data from source databases (**Figure 1b**). The CROssBARv2 data comprises around 2.7 million nodes spanning 14 distinct node types and approximately 12.6 million edges representing 51 different edge types (**Figure 1c**). To provide a structured and standardised representation of biological entities, we incorporated several ontologies, such as Gene Ontology and Mondo Disease Ontology. Additionally, we incorporated rich metadata as node and edge properties. The data processed by the adapter scripts is stored in the Neo4j graph database ^19^ to facilitate efficient querying, knowledge discovery, and in-depth analysis of complex biological relationships (e.g., protein-protein interactions, gene-disease associations, etc.), ensuring scalability and flexibility in complex network analyses (**Figure 1d**).

**Figure 1.**
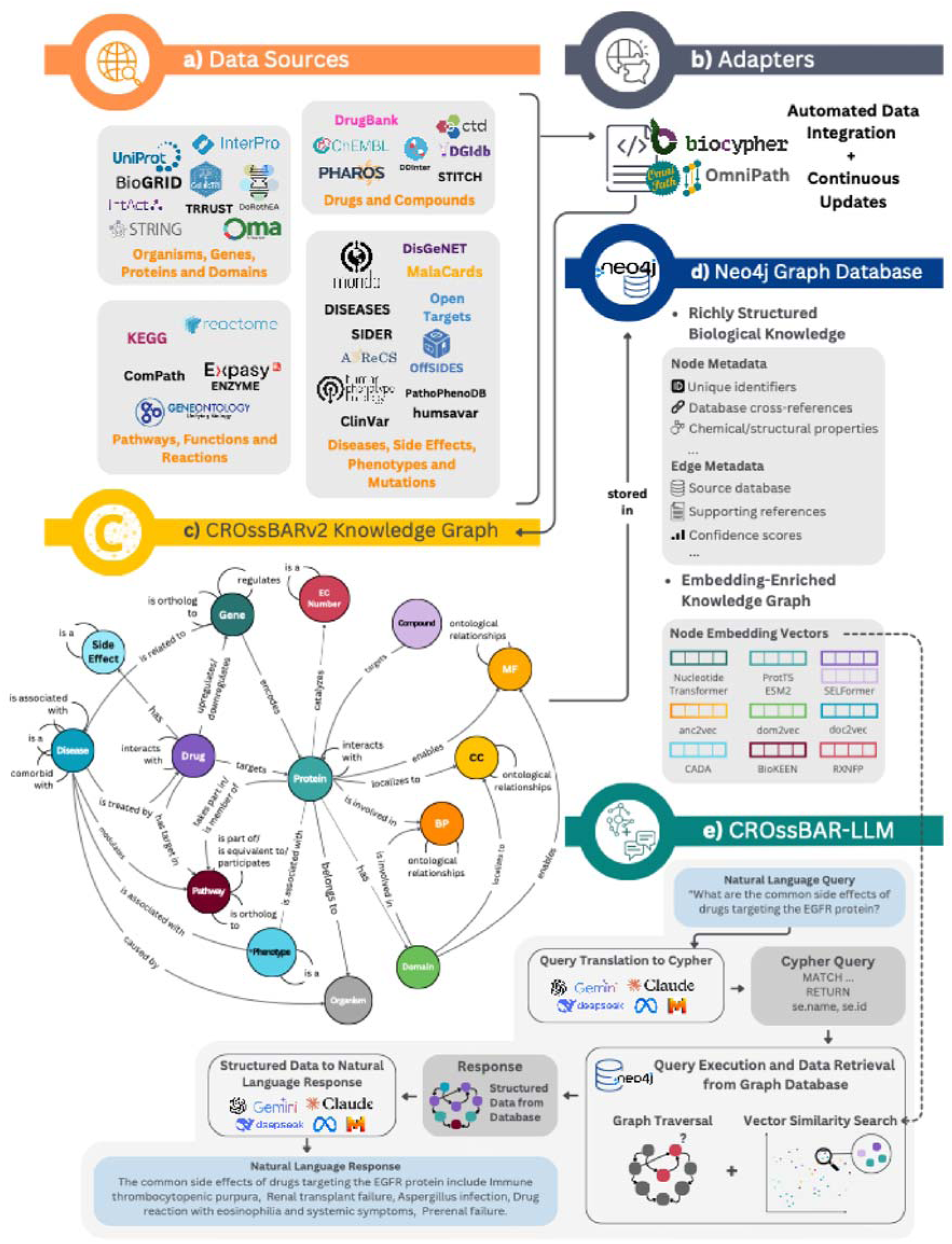
Overview of the CROssBARv2 workflow. **(a)** Integration of data from 34 well-established sources covering various biomedical domains. **(b)** Automatic retrieval, standardisation, and integration of source data using modular adapter scripts. **(c)** CROssBARv2 KG schema, comprising 14 node types and 51 edge types. **(d)** Storage of the KG in a Neo4j graph database, along with rich metadata and node embeddings computed using deep learning-based methods. **(e)** Execution of the CROssBAR-LLM workflow, which translates natural language queries into Cypher, executes the queries on KG, and synthesizes structured results into natural language responses; also supports vector-based similarity search.

Building upon this foundation, we further enhanced the semantic depth of CROssBARv2 by leveraging the vector index feature of the Neo4j graph database via generating and storing embeddings for biological entities in the KG, such as proteins, gene ontology terms, drugs and more (**Figure 1d**). These embeddings were generated using state-of-the-art deep representation learning models, such as ESM2 ^20^, ProtT5 ^21^, and SELFormer ^22^, which are trained to capture semantic information from amino acid sequences, atom/bond-centric notations, functional annotations, etc. (**Supplementary Table S5**). This vector-based representation enables sophisticated similarity searches, functional inference, and relationship predictions even when direct connections are absent in the KG.

To support large-scale data access, the CROssBARv2 platform also offers database dumps and the ability to build customised databases with configurable and reproducible adapter scripts, giving users the ability to access the entire KG or specific subsets as needed. Furthermore, we provide datasets formatted for direct use with popular deep learning frameworks, enabling researchers to incorporate the data into model training and testing workflows.

To enhance data extraction, exploration, and visualisation from the KG, we integrated three complementary layers on top of the graph database: (i) CROssBAR-LLM: a natural language question-answering system powered by LLMs that enables direct interaction with the KG (https://crossbarv2.hubiodatalab.com/llm) (**Figure 2a**), (ii) a GraphQL API for programmatic access allowing users to construct flexible, precisely defined queries, retrieving only the necessary information (https://crossbarv2.hubiodatalab.com/api), and (iii) the Neo4j Browser for interactive data exploration and visualisation (https://neo4j.crossbarv2.hubiodatalab.com/browser/?preselectAuthMethod=[NO_AUTH]&dbms=bolt://neo4j.crossbarv2.hubiodatalab.com) (**Figure 2c**). CROssBAR-LLM converts user questions into Cypher (Text-to-Cypher) queries that are automatically executed on the Neo4j database (https://crossbarv2.hubiodatalab.com/llm) (**Figure 1e**). The resulting structured output is then converted into coherent, contextual responses. This is facilitated via prompt engineering, enabling multiple LLM API calls at various stages of the pipeline (**Figure 1e**).

**Figure 2.**
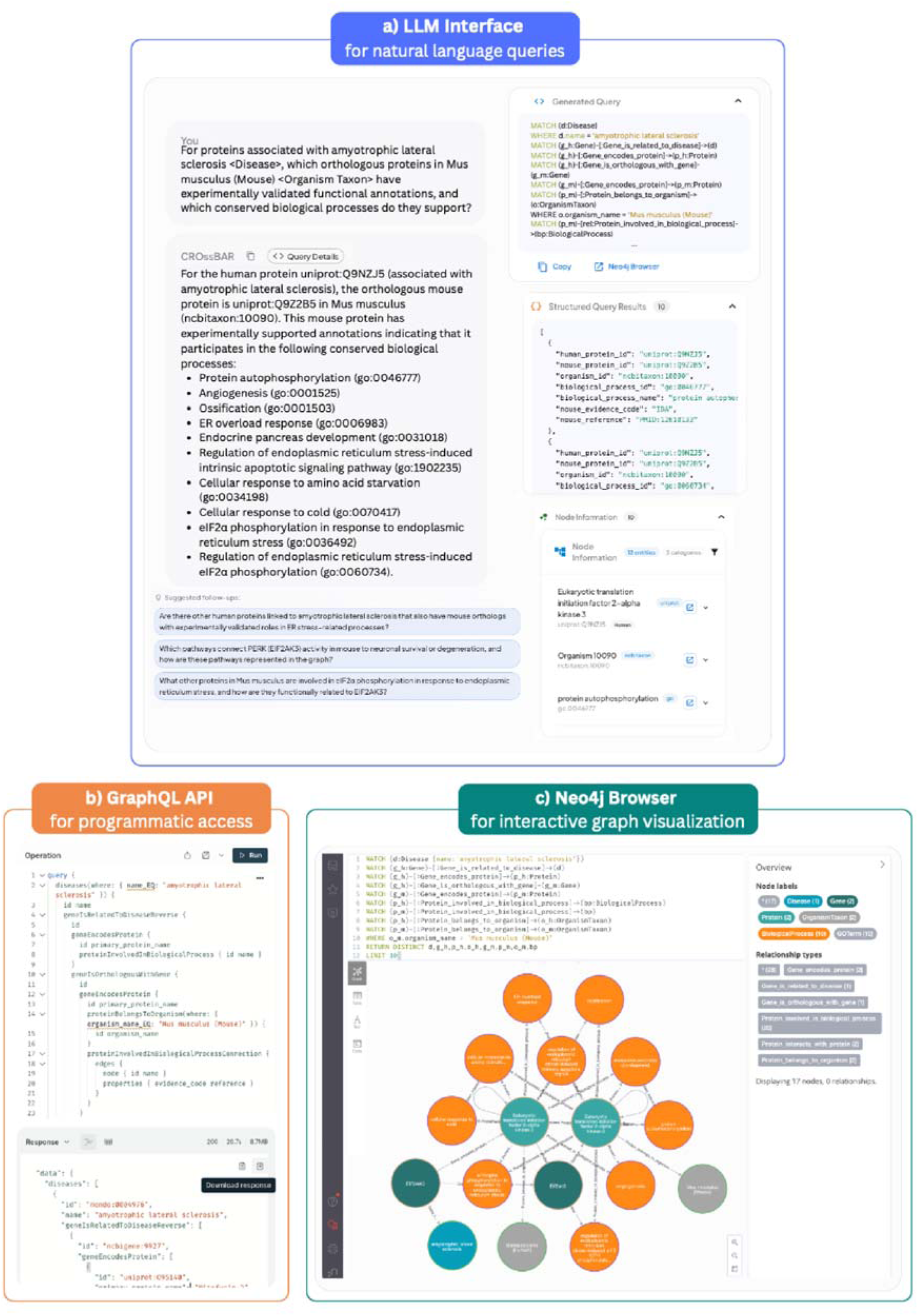
Exploration of the CROssBARv2 KG through three complementary interfaces. **a) Natural language querying via the CROssBAR-LLM tool** enables users to interact with the KG through questions written in plain language, supported by a backend that connects large language models with the Neo4j database (https://crossbarv2.hubiodatalab.com/llm). **b) Programmatic access through the GraphQL API** allows users to retrieve specific data and attributes, including nested relationships, through flexible and structured queries. **c) Interactive visual exploration via the Neo4j Browser** provides a graphical environment for navigating and inspecting nodes and relationships within the KG.

In the chapters that follow, we present (i) several use-case analyses that demonstrate the resource’s practical value, (ii) CROssBAR-LLM benchmarks for Cypher-query generation and biomedical question answering, accompanied by a qualitative comparison with standalone LLMs, (iii) a deep-learning framework that leverages CROssBARv2 to achieve state-of-the-art performance in predicting biological relationships, and (iv) a discussion of these findings alongside future perspectives.

### CROssBARv2 Knowledge Graph Use-Case Analyses

#### Part 1: Evaluating the biological validity of relationships integrated within the KG

To investigate the biological relevance of relationships within the KG, we applied the metapath2vec algorithm ^23^ to CROssBARv2. In contrast to the pretrained node embeddings employed in our system—which encode molecular or functional information derived from external sources (e.g., protein sequences or disease descriptions)—metapath2vec learns graph-structural representations by performing random walks over predefined metapaths, thereby rendering it particularly suitable for assessing relational patterns within a knowledge graph (KG). We applied metapath2vec in three exploratory use-cases focusing on protein, drug-disease, and pathway relationships, and analysed the resulting embedding spaces to evaluate whether biologically related or functionally similar nodes were positioned closer together. Such proximity would indicate that CROssBARv2’s relational structure itself encodes meaningful biological patterns that can support downstream tasks such as hypothesis generation, drug repurposing, functional annotation, and pathway analysis.

##### Exploring the functional significance of protein relationships

To test whether CROssBARv2 accurately represents functional protein relationships, we generated protein embeddings with metapath2vec using metapaths traversing protein-pathway, protein-GO term, and protein-EC number edges, such that each vector encodes participation in biological processes, molecular functions, and enzymatic reactions. In the t-SNE projection (**Figure 3a**) of obtained embedding space (dataset obtained from: ^24^), non-enzyme superfamilies (ion channels, membrane receptors, transporters, etc.) formed well-defined clusters, whereas enzyme families (hydrolases, proteases, transferases, etc.) were less distinct, but still reflected a broader separation. To quantify how much shared biology underlies these patterns, we counted shared edges: for every protein pair we recorded how many pathways, GO terms, or EC numbers they have in common. For each protein, we computed the mean number of unique shared edges with (i) proteins of the same family (intra-family) and (ii) proteins of all other families (inter-family), then averaged these values across family members. Non-enzyme families displayed ∼24-fold more intra-family than inter-family shared edges, while enzyme families showed an ∼8-fold enrichment (**Figure 3b**). The strong intra-family enrichment confirms that proximity in the embedding space is driven by genuine functional overlap captured through pathway, GO, and EC connections, demonstrating that the KG encodes biologically coherent protein family structure. To further contextualize these results, we compared obtained embeddings against Bioteque ^5^, a large-scale resource of pre-computed embeddings derived using metapath-guided node2vec over a KG integrating more than 150 biomedical datasets. Applying agglomerative clustering, CROssBARv2 achieved purity scores of 0.504 for enzyme families and 0.818 for non-enzymes, compared to 0.445 and 0.773 for Bioteque, respectively, confirming that our KG captures functional structure with enhanced biological coherence. Details of this analysis and its results can be found in **Supplementary Information S2.1**.

**Figure 3.**
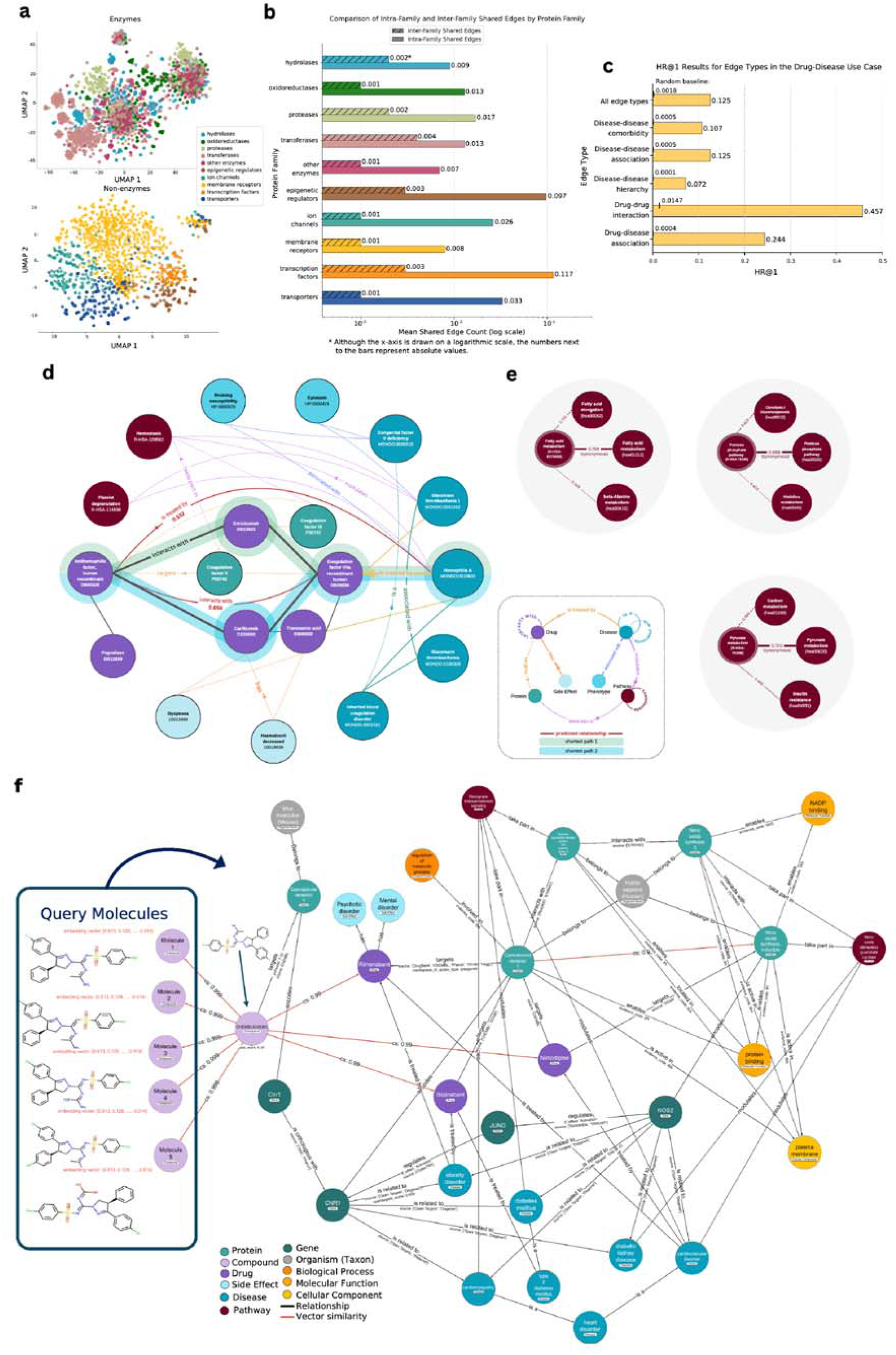
CROssBARv2 use-case analyses. <u>Part 1</u>: Metapath2vec centred case study on assessing biological relevance across protein, drug-disease, and pathway interactions within the CROssBARv2 KG. (**a)** t-SNE projections of the metapath2vec embedding space for protein nodes. Two separate projections were made for enzymes and non-enzyme proteins. (**b)** Comparison of average in-family and cross-family shared edge counts among protein families in the protein use-case. The x-axis is log-scaled, and raw counts are displayed to the right of each bar. **(c)** Hit rate at 1 (HR@1) values for edge types in the drug-disease association use-case. HR@1 evaluates the model’s ability to place a true relational neighbour at the top of the ranking based on path-based similarity. Baseline HR@1 values are shown above each bar as reference markers. **(d)** Key connections among selected nodes, including Antihemophilic Factor, human recombinant (DB00025), Coagulation factor VIIa Recombinant Human (DB00036), Hemophilia A (MONDO:0010602) in the KG. Predicted relationships based on path-based similarity and supported by literature between DB00025 and DB00036,as well as DB00025 and MONDO:0010602, are indicated by red edge lines, with their cosine similarities displayed below the edge labels. The two shortest paths between DB00025 and MONDO:0010602 are highlighted with coloured backgrounds. **(e)** Pathway synonymy relationships and metapath2vec embedding similarity scores between selected nodes in the pathway use-case. Solid lines represent known synonymy between pathways, while dotted lines connect pathways without a synonymy relationship. Each line displays the embedding similarity score for the node pairs, with thicker lines denoting higher similarity scores. **(f) <u>Part 2</u>: Subgraph extracted from CROssBARv2 KG for the elucidation of potential therapeutic mechanisms of de novo molecules** (node colors denote entity types; black edges indicate curated KG relationships; red edges show vector similarity). De novo query molecules (left) were embedded using SELFormer and searched against the CROssBARv2 KG via vector similarity search, identifying CHEMBL4076361 (cosine similarity > 0.99). The resulting subgraph (right) links CHEMBL4076361 to mouse Cnr1 and further to human CNR1 through orthology relationships. CNR1 and NOS2 proteins/genes have shared biological context including disease associations (e.g., cardiovascular disorders, diabetes mellitus), molecular functions (protein binding), subcellular localisation (plasma membrane), transcriptional activation (mediated by JUND). The shortest path length between CNR1 and NOS2 proteins in the KG is 3; one such path, proceeding via Guanine nucleotide-binding protein G(i) subunit alpha-2 and Nitric oxide synthase 3, is depicted in the graph (right upper). CNR1 is involved in regulation of metabolic process and takes part in Retrograde Endocannabinoid Signaling pathway, while NOS2 protein enables NADP binding and participates in Nitric Oxide Stimulates Guanylate Cyclase pathway. Notably, CHEMBL4076361 showed significant cosine similarity (>0.99) to Rimonabant, a once-approved anti-obesity drug and the inverse agonist of the Cannabinoid receptor 1 that was withdrawn from clinical use due to neuropsychiatric side effects, also depicted in the graph. This example illustrates CROssBARv2’s ability to infer potential mechanisms of new compounds absent from existing biomedical resources.

##### Analysing drug-disease associations for therapeutic relevance

We evaluated the KG’s potential for modelling drug and disease relationships by testing whether the proximity of drug and disease node embeddings generated with metapath2vec reflects their true biological associations, as captured by disease-drug associations, drug-drug interactions, and three disease-disease relations (hierarchy, association, comorbidity). We used hit ratio at 1 (HR@1) to measure how often a node’s closest neighbour in the embedding space is linked by a true KG edge, treating nearest-neighbour retrieval as edge prediction. As a baseline, we use the likelihood that a randomly selected candidate corresponds to a true relation, determined by the node’s ground-truth edge count divided by the size of the candidate space. The model recovered 45.7% of known drug-drug interaction edges (HR@1 = 0.457, **Figure 3c**) and achieved lower but meaningful scores for the other classes, confirming that heterogeneous graph information supports relationship recovery. When compared with the HR@1 baselines, the model shows clear gains across all edge types. Coagulation factor VIIa (DB00036) exemplifies this robustness, being positioned near its annotated diseases Glanzmann thrombasthenia 1 and haemophilia A, and with five interacting drugs, validating embedding reliability (**Figure 3d**). Anti-haemophilic factor (DB00025), the next-closest drug, lacks an explicit interaction edge yet is linked to DB00036 through short paths and, while not annotated for haemophilia A in source databases, is described as such in DrugBank and the literature ^25,26^, illustrating the method’s ability to suggest novel associations (**Figure 3d**). While pathway, side-effect and phenotype nodes were not included in metapaths, **Figure 3d** shows they form indirect links between DB00025 and haemophilia A. Incorporating them could enrich embeddings and improve the system’s ability to identify novel therapeutic associations. Details of this analysis and its results can be found in **Supplementary Information S2.2**.

##### Assessing cross-database pathway synonymy

Pathway embeddings, generated with metapath2vec from hierarchical pathway-pathway and protein-pathway relations, were evaluated by checking whether curated KEGG-Reactome synonym pairs appeared close together in the vector space (**Figure 3e**). Across all pairs, cosine-similarity rankings yielded a hit ratio at 10 of 0.426, i.e. 43% of synonym pairs appear within the ten most-similar neighbours despite the benchmark accepting only exact name matches. To illustrate this biological coherence with a concrete example, the Reactome fatty-acid metabolism pathway R-HSA-8978868 is close to its KEGG synonym hsa01212 (similarity 0.759) and to the related elongation pathway hsa00062 (0.735), whereas an unrelated β-alanine pathway (hsa00410) scores lower (0.549). These results show that integrating pathway hierarchies and protein associations enables embeddings to recover biologically meaningful pathway relationships. Details of this analysis and its results can be found in **Supplementary Information S2.3**.

#### Part 2: Elucidating Potential Therapeutic Mechanisms of De Novo Molecules via CROssBARv2’s Hybrid Approach

To evaluate the effectiveness of CROssBARv2’s hybrid search and the breadth of its integrated knowledge, we considered a drug-discovery scenario based on a series of de novo, drug-like small molecules. Specifically, we defined one drug-like compound and generated five closely related virtual analogues (**Figure 3f**, left), all of which are absent from public chemistry resources and from CROssBARv2. This setup mimics an expert computational chemist exploring a new compound series and is intended to illustrate how CROssBARv2 can consolidate and surface relevant biomedical evidence.

For each query molecule, we performed a vector-similarity search over the KG using SELFormer-derived embeddings ^22^, which are already integrated into CROssBARv2 for compound nodes. In all cases, the search converged on the same proxy molecule, CHEMBL4076361, with cosine similarity >0.99 to all five query structures, indicating that the model reliably identifies close structural neighbours. We then expanded an automated contextual subgraph around CHEMBL4076361, capturing shortest paths, interaction types and supporting annotations to reveal its biological neighbourhood (**Figure 3f**, right).

The resulting subgraph showed that CHEMBL4076361 targets mouse Cnr1, which maps via orthology to the human CNR1 gene. Further exploration identified NOS2 as a protein with high vector similarity (0.93) and overlapping biological context with CNR1, including shared plasma-membrane localization and protein-binding activity. Both genes were associated with metabolic and cardiovascular conditions such as diabetes and cardiovascular disease, supported by multiple studies ^27–34^. In addition, anti-obesity agents Ibipinabant and Rimonabant, which target CNR1, and the cardiovascular drug Nimodipine, for which the KG reports interactions with both CNR1 and NOS2, exhibited a cosine similarity of ≈0.99 to CHEMBL4076361. Together, these connections suggest that the de novo molecules may plausibly act as dual modulators of CNR1 and NOS2, with potential relevance for metabolic and cardiovascular indications. These hypotheses will require further computational and experimental validation; however, CROssBARv2 assembles a coherent mechanistic and disease context that can help prioritise such hypotheses and reduce the effort and time between in silico exploration and downstream wet-lab experimentation.

This analysis was conducted in a “blinded” manner. We first selected a drug-like compound with experimental bioactivity data and generated the five de novo analogues by introducing minor atom-level substituent and functional-group changes while preserving the core scaffold. Upon unblinding, we confirmed that the parent molecule in this series is MRI-1569, a dual inhibitor of CNR1 and NOS2 ^35^. The biological context recovered by CROssBARv2—CNR1/NOS2 involvement, metabolic and cardiovascular indications, and similarity to Ibipinabant, Rimonabant and Nimodipine—recapitulates the known pharmacology of MRI-1569 ^35^; ^36^. Thus, starting from virtual molecules that do not exist in any database, CROssBARv2 reconstructs a literature consistent mechanistic and disease profile via combined semantic similarity search and graph traversal. Details of this analysis are provided in **Supplementary Information S2.4**.

### CROssBAR-LLM

Large Language Models (LLMs) frequently suffer from hallucinations, generating plausible but inaccurate or unsupported responses, particularly problematic in technical contexts such as complex scientific question answering, where precision and factual correctness are crucial. Utilising highly structured and semantically rich relationship-based information, like the CROssBARv2 biomedical knowledge graph (KG), for scientific queries can mitigate this issue by providing explicit, verifiable connections between entities and concepts. However, KGs are difficult to communicate with, usually requiring long and complex database queries written in specialised programming languages. Similarly, digesting the structured output of these queries requires additional processing steps and expertise. Integrating biomedical knowledge graphs with LLMs can offer a robust solution to both scientific accuracy and practical usability challenges, effectively combining the structured, high-fidelity scientific content inherent in knowledge graphs with the intuitive, natural language querying capabilities of LLMs.

To address these limitations, we propose CROssBAR-LLM (Figure 2a), an intuitive online system that integrates CROssBARv2 KG with LLMs for practical, factual, and natural language-based scientific question answering (https://crossbarv2.hubiodatalab.com/llm). As illustrated in **Figure 1e**, the process begins with the user’s biomedical query, posed to CROssBAR-LLM in natural language. The underlying LLM generates a corresponding graph database query designed to retrieve relevant information from the KG. This query is then executed on the KG at the backend, extracting structured output data that captures detailed relationships and attributes among biological entities. The retrieved structured data is subsequently returned to the LLM, which synthesises it into a coherent natural language response. Obtaining plausible results from this process required complex prompt engineering and a high volume of testing (please see Methods for details).

In addition to direct graph-based querying, users can optionally enable vector-based semantic similarity search. When activated, the system leverages precomputed embeddings to identify biologically related entities beyond exact graph matches, allowing retrieval of not only explicitly connected nodes but also semantically similar entities that may support broader hypothesis generation. The interface supports iterative analysis: users can continue the interaction by submitting their own follow-up questions or by selecting follow-up suggestions proposed by the LLM. This conversational workflow enables progressive query refinement and deeper exploration of the knowledge graph. Example queries for both functionalities, along with relevant information, are provided in **Figure S1** and **Supplementary Information S3**.

#### A qualitative comparison between CROssBAR-LLM and standalone LLMs in complex biomedical question-answering

To investigate the potential of CROssBAR-LLM for improving accuracy, contextual relevance, and overall quality of biomedical question answering, we conducted a qualitative use-case study. Here, we compared the responses generated by the standalone LLM with those produced by CROssBAR-LLM. The queries were intentionally designed as complex, multi-entity scenarios to probe the limits of CROssBAR-LLM’s ability to integrate diverse biomedical entities and relationships in challenging settings, rather than to reflect real user-generated research questions. For these comparisons, we selected GPT-4o, a state-of-the-art LLM at the time of writing, recognized for its robust natural language processing capabilities and ranking among the top performers in our LLM benchmark (please see “CROssBAR-LLM benchmarks on Cypher query generation and biomedical question answering”). GPT-4o was specifically prompted to answer the queries using its web search functionality, ensuring access to the most recent and relevant information available. The two selected queries and their respective analyses are discussed in detail below.

As our first complex query—“*What drugs are used for ‘obesity disorder’ disease and have drug interactions with ‘Ceritinib’, while also targeting proteins involved in the ‘ABC transporters’ pathway?*”—was issued to evaluate multi-layer reasoning. Correct answers must reconcile (i) drug-disease indications, (ii) drug-drug interactions with the tyrosine-kinase inhibitor ceritinib (mediated mainly through CYP3A4 metabolism or transporter-related QT-prolongation effects), and (iii) drug-protein links involving ABC transporters such as ABCB1 that regulate intracellular drug concentrations. Running this query through CROssBAR-LLM yielded twelve qualifying drugs (**Supplementary Table S1; Figure 4a**). Among them, sertraline (eating-disorder therapy), rosuvastatin (dyslipidaemia), and bromocriptine (type-2 diabetes) possess strong obesity-related indications and preserve all three required relationship types: they are associated with obesity or its comorbidities, documented to interact with ceritinib via CYP3A4 or transporter mechanisms, and bind proteins that participate in the ABC-transporter pathway. Other returned agents (e.g., sevoflurane, propofol, azithromycin, hydrocortisone) appear because their broad clinical use generates multiple disease annotations, drug-interaction records with ceritinib, and transporter involvement. Querying less clearly defined disease entities such as “obesity disorder” can inevitably introduce misleading results due to broad or indirect annotations. Our rationale for including such an example was to illustrate that CROssBAR-LLM is still able to generate interpretable candidate answers in ambiguous scenarios where standalone LLMs typically fail.

**Figure 4.**
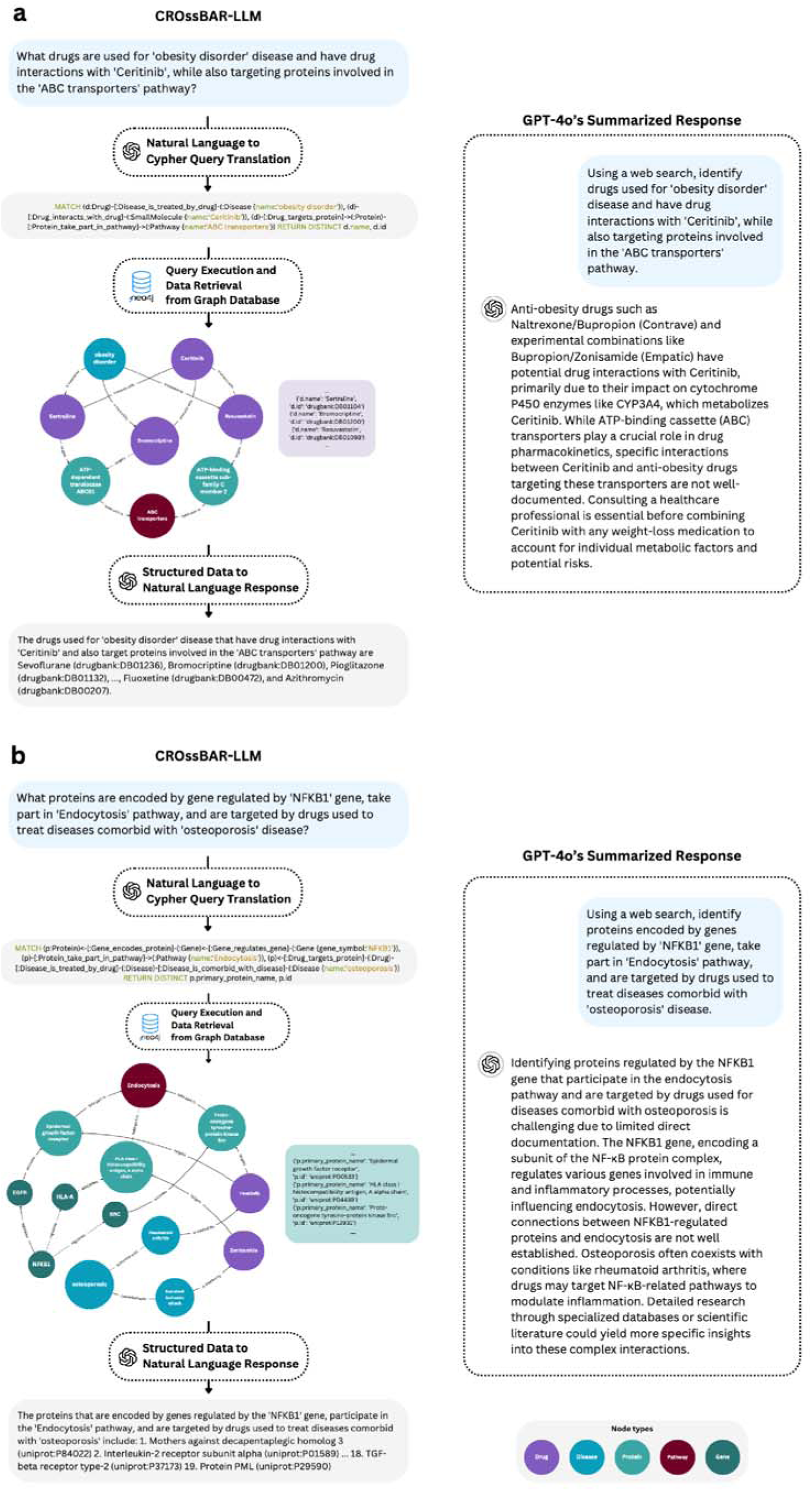
Comparison of responses generated by the CROssBAR-LLM system and a standalone LLM (GPT-4o) for two complex biomedical queries. For each query, a subgraph of the retrieved results is visualised, highlighting key entities and interactions in CROssBARv2 KG. The natural language response of both CROssBAR-LLM and GPT-4o are shown along with each other. **(a)** The query on identifying drugs used for obesity that interact with Ceritinib and target proteins involved in the ABC transporters pathway. The displayed subgraph features three selected key drugs—Sertraline, Bromocriptine, and Rosuvastatin—from the system’s response. **(b)** The query on identifying proteins regulated by NFKB1, involved in the endocytosis pathway, and targeted by drugs used to treat diseases comorbid with osteoporosis. The displayed subgraph features three selected key proteins—Epidermal growth factor receptor, HLA class I histocompatibility antigen A alpha chain, and Proto-oncogene tyrosine-protein kinase Src—from the system’s response.

When the same query was posed to GPT-4o using its web-search mode, the model produced a narrower answer: naltrexone/bupropion and bupropion/zonisamide combination therapies, citing ceritinib’s CYP3A4 metabolism but admitting that specific ABC-transporter interactions are “poorly documented” (**Figure 4a**). GPT-4o therefore met only part of the specification, lacking explicit transporter-level evidence and omitting other valid candidates. On the other hand, CROssBAR-LLM integrated drug-disease, pharmacokinetic and pathway knowledge to recover a more complete and mechanistically grounded drug set, demonstrating superior capability for answering biologically intricate queries. Details of this analysis and its results, including the full responses obtained from both LLMs, can be found in **Supplementary Information S4.1**.

Our second query: “*What proteins are encoded by gene regulated by ‘NFKB1’ gene, take part in ‘Endocytosis’ pathway, and are targeted by drugs used to treat diseases comorbid with ‘osteoporosis’ disease?*” was formulated as a highly complex example, combining multiple concurrent requirements to assess CROssBAR-LLM’s ability to perform deep, integrative reasoning across diverse information layers and the extent to which such queries can be meaningfully addressed. CROssBAR-LLM executed this multi-step integration by (i) extracting genes with regulated by NFKB1, (ii) intersecting their protein products with the KEGG Endocytosis pathway (hsa04144), and (iii) filtering for proteins bound by drugs whose indications overlap diseases comorbid with osteoporosis (e.g., arthritis, depression, cardiovascular disease). The pipeline returned 19 proteins (**Supplementary Table S2**) that satisfy all constraints. High-significance examples include proto-oncogene SRC (UniProt ID: P12931), which modulates survival and migration signalling, three HLA class I molecules (HLA-G, -E, -A) central to antigen presentation, and Epidermal growth factor receptor (P00533), a driver of proliferation via growth-factor cascades. **Figure 4b** depicts their tripartite relationships: regulatory links from NFKB1, pathway membership edges to Endocytosis, and drug-target interactions with agents such as imatinib and zonisamide—therapeutics approved for comorbid conditions. Additional output proteins contribute to immune modulation, cytoskeletal dynamics and vesicular trafficking, underscoring the biological coherence of the set.

When the same query was posed to GPT-4o with web-search enabled, the model provided background for each individual aspect of the query but failed to identify proteins fulfilling all criteria together, finally, recommending manual literature searches (**Figure 4b**). Its narrative lacked traceable source links and evidence, illustrating the limitations of free-text reasoning without an underlying structured KG. In contrast, CROssBAR-LLM produced a complete and source-traceable answer, highlighting the advantage of coupling LLMs with curated heterogeneous graphs for complex biomedical inquiry. Detailed results and discussion of query example 2 can be found in **Supplementary Information S4.2**.

#### CROssBAR-LLM benchmarks on Cypher query generation and biomedical question answering

We examined LLM backbones used in CROssBAR-LLM with two objectives: (i) to quantify their ability to translate natural-language queries into valid Cypher—essential for interacting with the Neo4j graph database that hosts CROssBARv2—and (ii) to determine whether augmenting these models with the CROssBARv2 KG improves biomedical question answering, both on an internal benchmark and the independent GeneTuring ^37^ benchmark.

To evaluate the capability of CROssBAR-LLM, backed by different LLMs, in generating valid Cypher queries, we presented 50 curated questions (e.g., “Find all drugs that decrease the expression of genes related to ‘Duchenne and Becker muscular dystrophy’.”) to each model. The performance results in **Figure 5a** revealed that open-source models, including DeepSeek R1 (41/50), LLaMA 3.1-405B (39/50) and Mistral Large 2 (41/50), demonstrated performance on par with proprietary models (e.g., Claude 3.5 and GPT-4 series). To specifically assess performance on complex graph traversals, we analyzed a subset of 11 questions requiring queries with a path length of 3 or more hops (e.g., “Which side effects are caused by drugs that target proteins involved in the ‘chemical synaptic transmission’ biological process?”). On this challenging subset, leading models like the GPT-4 series and DeepSeek R1 performed strongly (9/11), while smaller models like Llama 3 70B and GPT-4o-mini struggled (2/11). These results are consistent with the full 50-question benchmark, where larger, more advanced models consistently outperformed smaller ones.

**Figure 5.**
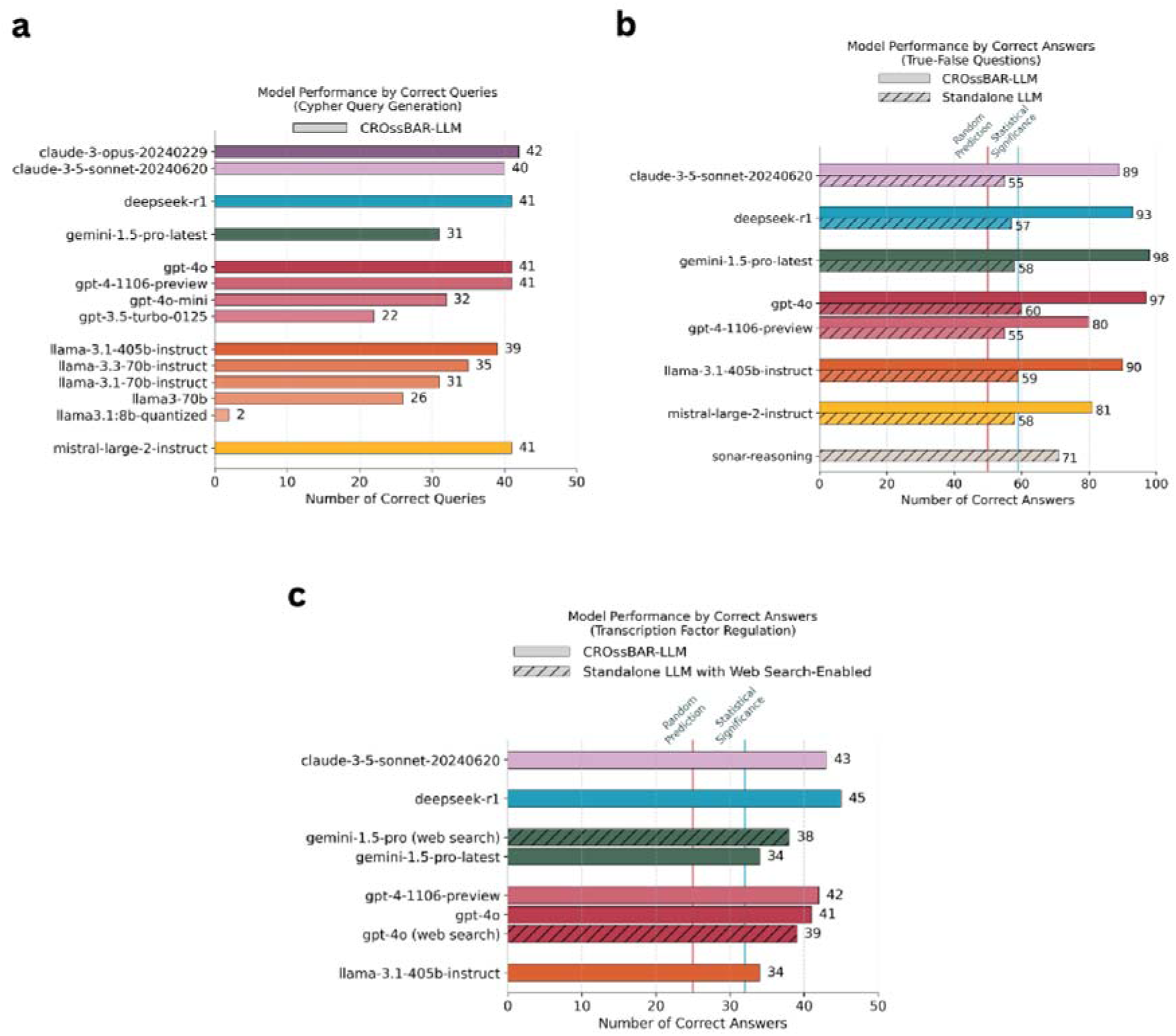
CROssBAR-LLM benchmarking performance results on Cypher query generation and biomedical question answering tasks. **(a)** The number of correct Cypher queries produced by various LLMs utilised in CROssBAR-LLM (out of 50 cases). **(b)** The numbers of correct answers for standalone LLMs (striped bars) and CROssBAR-LLMs (solid bars) when answering in-house curated “true-false” based biomedical questions (out of 100 cases). **(c)** The number of correct answers achieved by CROssBAR-LLMs (solid bars) compared to web search-augmented LLMs (striped bars) on the “transcription factor regulation” module of the GeneTuring benchmark (out of 50 cases). For b and c, the random prediction line (red) indicates the expected performance of a random predictor at 50%. The statistical significance line (blue) indicates models performing significantly better than random prediction.

In our knowledge-augmented biomedical question answering task, we employed two datasets to measure the impact of coupling LLMs with the CROssBARv2 KG. The first consisted 100 internally curated true/false questions and the second comprised 50 binary choice questions as activation/repression “transcription factor regulation” module of externally curated GeneTuring benchmark. On the internal true/false benchmark (e.g., “AGER Gene is associated with acute monocytic leukemia. Answer only True or False”), we compared CROssBAR-LLMs with standalone LLMs and a web-search augmented model, Sonar Reasoning ^38^. Interestingly, most standalone LLMs do not perform significantly better than random guessing. Conversely, augmenting LLMs with the CROssBARv2 KG dramatically boosted performance, enabling near-perfect accuracy from models like Gemini 1.5 Pro (98%) and GPT-4o (97%), far surpassing both standalone LLMs and the Sonar Reasoning (71%) (**Figure 5b**).

Performance trends persisted on the GeneTuring set (e.g., “Does transcription factor MSC activate or repress gene CDC6?”). Our evaluation demonstrated that CROssBAR-LLMs generally outperformed web-search augmented LLMs (**Figure 5c**). Claude 3.5 Sonnet and DeepSeek R1 backed CROssBAR-LLMs achieved the highest accuracy (43/50 and 45/50, respectively), whereas web search-augmented standalone GPT-4o and Gemini-1.5-Pro performed slightly worse (39/50 and 38/50, respectively). The full question and answer sets are available in the **Supplementary Files**, and additional findings are reported in the **Supplementary Information S5**.

Overall, CROssBAR-LLM demonstrated strong proficiency in natural language understanding and precise translation into Cypher queries, even for complex multi-hop graph traversals. Furthermore, CROssBAR-LLM, which leverages our systematically organized KG, consistently outperformed both standalone and web search-augmented LLMs in biomedical question answering, underscoring the importance of structured biomedical knowledge in improving LLM performance in specialised domains.

### Deep learning-based prediction of biological relationships using CROssBARv2

To demonstrate the applicability of the CROssBARv2 KG for deep learning-based prediction of biological relationships, we trained protein function prediction models. Protein function annotation remains a fundamental challenge in molecular biology; heterogeneous KGs can address it by unifying sequence, structural, pathway and interaction data. For this, we introduce ProtHGT, a Heterogeneous Graph Transformer (HGT)-based model that learns on the CROssBARv2 KG and treats protein-Gene Ontology (GO) links as edges to be predicted^39^. Initial node features (sequence, domains, pathways, etc.) are refined through HGT’s type-aware message passing, which uses independent weight matrices for each node and edge type to learn rich, context-specific representations. The final protein and GO embeddings are concatenated and scored by a two-layer MLP with sigmoid output. Each HGT block (**Figure 6a**) consists of three key submodules and an attention mechanism that determines the influence of each source node on a target node using type-specific projections; a message-passing module that transforms source node information through edge-type-specific operations; and a feature aggregation step where messages are combined using attention weights, followed by non-linear transformation, a residual connection, and a final type-specific projection to update the target node’s representation. By stacking multiple HGT blocks, the model enables information to propagate throughout the graph, allowing nodes to capture both local and long-range dependencies.

**Figure 6.**
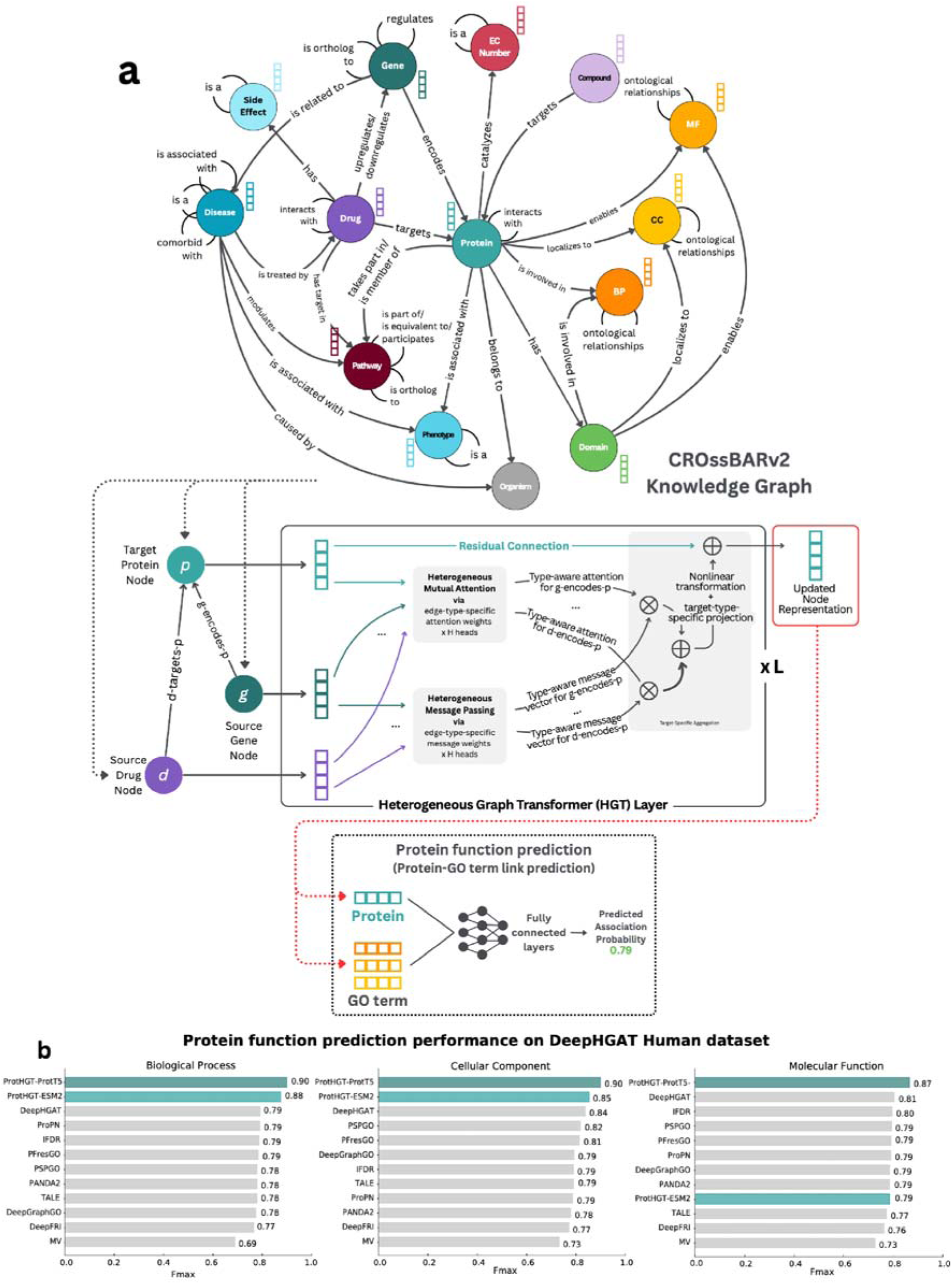
Deep learning-based utilization of CROssBARv2 for biological relationship prediction. **(a)** Overview of Heterogeneous Graph Transformer (HGT) based ProtHGT model and its training on the CROssBARv2 knowledge graph for protein function prediction. **(b)** Protein function prediction performance evaluated on the DeepHGAT Human dataset using Fmax scores. ProtHGT models that utilise ESM2 and ProtT5 protein language models achieved top performance in all categories.

Separate models were trained for the molecular-function, cellular-component and biological-process sub-ontologies. To ensure time-aware evaluation, we built a separate date-restricted version of the KG by removing all protein-GO edges and reinserting only those available during the training period of the DeepHGAT (≤ Sep 2021) ^40^ benchmark; annotations added in the subsequent test windows served as ground truth. In the DeepHGAT benchmark, ProtHGT achieved the highest Fmax across all three sub-ontologies (**Figure 6b**), outperforming heterogeneous-graph competitor DeepHGAT, PPI-based PSPGO, and state-of-the-art sequence models. The improvement stems from CROssBARv2’s broader relational context—including pathways, domains, and ontological edges—absent from models restricted to sequences or single network types.

Because ProtHGT operates on an explicit multi-relational KG, each prediction can be traced through specific pathway, domain, or interaction paths, providing mechanistic insight that most sequence-only approaches lack. These results confirm that deep learning on a richly integrated KG yields both state-of-the-art accuracy and interpretable protein function predictions. Details of this analysis and its results can be found in **Supplementary Information S6**.

## Discussion & Conclusion

CROssBARv2 effectively addresses the major challenges in existing KG platforms by providing a comprehensive, flexible, metadata-rich, and user-friendly solution for biomedical data integration. By systematically unifying heterogeneous biological data from over 34 sources into a KG, CROssBARv2 enables researchers to explore complex biomedical relationships with unprecedented depth and precision. The system introduces an automated and configurable data integration pipeline via “adapter” scripts, ensuring long-term maintainability and reproducibility. To bridge the gap between technical complexity and user accessibility, the system offers a multi-layered interface, including a GraphQL API for programmatic access, an interactive Neo4j Browser for visual exploration, and an LLM-powered natural language question answering tool. This design ensures that both computational and experimental researchers can efficiently query and analyse the KG without requiring specialized graph database expertise.

CROssBARv2 uses Neo4j’s vector indexing combined with deep representation learning models like ESM2, ProtT5, and SELFormer to create embeddings that capture key semantic features of biological entities, enriching the KG. Vector similarity search extends the capabilities of CROssBARv2 by identifying entities with shared functional, structural, or semantic properties (e.g., proteins with analogous enzymatic domains or drugs with similar pharmacophores), even without direct connections in the KG. Moreover, the use of embeddings ensures compatibility with machine learning frameworks, providing feature-rich inputs for advanced computational models, such as deep graph neural networks. The synergistic combination of graph traversal and vector search – a hybrid search approach – provides a particularly powerful toolset, allowing for multifaceted queries that leverage both explicit graph connections and implicit semantic similarities. For instance, users can start with a known disease-associated gene (via graph traversal) and then identify structurally similar proteins (via vector search) to uncover novel targets or repurposable drugs. The use-case study of de novo small molecules demonstrates this capability, where CROssBARv2 successfully mapped uncharacterized molecules to existing biomedical knowledge, despite these novel structures being absent from chemical databases.

In our tests, augmenting LLMs with the structured biomedical knowledge within CROssBARv2 led to substantial improvements in biomedical question-answering accuracy compared to standalone models and even web search strategies, underscoring the value of curated, domain-specific knowledge. Rigorous analysis using metapath2vec embeddings validated the KG’s ability to capture meaningful relationships, and use-cases demonstrated its versatility across complex biological tasks, including drug repurposing, pathway analysis, and functional annotation. Additionally, deep learning models trained on CROssBARv2 achieved state-of-the-art performance in protein function prediction, confirming that its heterogeneous structure enables precise and context-rich representation learning. Beyond predictive accuracy, CROssBARv2 also offers interpretability, enabling predictions to be traced to underlying biological relationships from which they arise. Collectively, these features address longstanding challenges in usability, reproducibility, adaptability, and explainability, positioning CROssBARv2 as a powerful framework and versatile tool for hypothesis generation in systems biology, drug discovery, and precision medicine.

The databases used to construct biomedical KGs contain data ranging from experimentally validated findings to automatically extracted information from text mining, each with varying degrees of reliability. This heterogeneity and inherent uncertainty often reduce the reliability and credibility of edges within biomedical KGs. To address this challenge, we enriched the CROssBARv2 KG with comprehensive metadata as node and edge properties, including source databases, supporting references (e.g., PubMed identifiers, evidence codes), and confidence scores. This metadata empowers researchers to implement diverse filtering strategies during knowledge discovery, such as prioritizing protein-protein interactions reported in multiple databases or those with high confidence scores, thereby focusing their analyses on the most reliable biological relationships. Such enhancements not only facilitate well-founded information retrieval and provenance tracking but also improve the overall reliability and interpretability of the CROssBARv2 system. To further enhance knowledge discovery, CROssBARv2 integrates embeddings from diverse deep learning models, enabling cross-domain semantic similarity searches beyond traditional protein-centric tasks. A detailed discussion of these aspects is provided in the **Supplementary Information S7.1**.

With CROssBARv2, we aim to consolidate fragmented biomedical data into comprehensive, interoperable, and sustainable KG. To accomplish this, CROssBARv2 utilises cross-references between identifiers of databases and ontologies to integrate equivalent or related biological entities from diverse sources into standardised node and edge representations. However, according to our observations, this approach faces two key challenges: (i) incomplete or inconsistent cross-referencing across sources, and (ii) managing the diverse conceptual granularity across different disease classification systems. We elaborate on these issues in **Supplementary Information S7.2**.

With their growing sophistication and vast internal knowledge, LLMs can be perceived as potential replacements for knowledge bases. In this context, we believe that CROssBAR and similarly rich biomedical data sources will remain highly relevant. LLM outputs, while steadily improving, are often prone to hallucination, lack grounding in real-time data, and underperform on domain-specific tasks without external tool use ^41,42^ as a result, they require trustworthy external knowledge sources. One way to supply that knowledge is through Model Context Protocol (MCP)^43^: the LLM issues a structured API request that triggers an external operation—e.g., a database query, a computation service, or laboratory equipment control—and then integrates the returned result into its reasoning chain. These calls ground the model in up=to=date, domain=specific data, reduce hallucination, and ensure that outputs meet biomedical accuracy and regulatory standards. This is particularly critical in biomedical applications, where accurate, interpretable results are essential for downstream decision-making. For instance, biomedical AI agents use LLMs alongside external tools to access curated databases, control lab hardware, and iteratively refine hypotheses based on feedback ^44^. In this setting, CROssBARv2’s unified KG of curated biomolecular relationships can be very valuable. By interacting with the CROssBARv2 KG, an LLM can retrieve verified relational and contextualized information on demand, reason over it, and produce answers that remain interpretable and auditable. Thus, rather than replacing resources like CROssBAR, increasingly sophisticated LLMs are likely to continue benefiting from them, both for grounding and for systematic evaluation, to deliver more reliable, actionable biomedical insights. In addition to these considerations, it is also important to consider how different KG-LLM integration strategies compare in practice. In **Supplementary Information S7.3**, we contrast our Text-to-Cypher approach with retrieval-augmented generation (RAG)-based alternatives.

Future releases of CROssBAR will first expand on multi-omics—transcriptomics (e.g., Expression Atlas ^45^, GTEx ^46^), proteomics (PaxDb ^47^, ProteomicsDB ^48^), metabolomics (Metabolomics Workbench ^49^, HMDB ^50^), epigenomics (ENCODE ^51^, Roadmap Epigenomics ^52^, MethBank ^53^), single-cell omics (CELLxGENE ^54^, Single Cell Expression Atlas), and microbiomics/metagenomics (MGnify ^55^)—to support cross-scale reasoning from genotype to phenotype. In parallel, building on the existing shared components between CROssBARv2 and OmniPath^56,57^, we will develop a tighter integration of database building pipelines for improved reproducibility and the incorporation of more resources in CROssBAR. As OmniPath expands, so will CROssBAR, in particular adding nutrition-focused resources such as FooDB ^58^ and FoodOn ^59^ linking diet, metabolism, and disease. Semantic depth will be strengthened with embeddings from state-of-the-art representation models such as ESM3 ^60^, Evo 2 ^61^, and Uni-Mol models ^62,63^, and forthcoming structure-aware multi-modal chemical and biological representation learning models. On the CROssBAR-LLM front, we are exploring the evolution of CROssBAR-LLM toward an multi-agentic system that maintains conversational context and autonomously refines query generation, and we are considering adopting MCP to standardise and streamline LLM interactions with the CROssBARv2 KG. Furthermore, we are implementing a GraphRAG-based service tailored for KGs, inspired by approaches such as SimGRAG ^64^, which is currently undergoing internal testing and is planned to be released shortly to further enhance biomedical question answering.

Overall, CROssBARv2 helps consolidate fragmented biomedical sources into a maintainable, provenance-rich knowledge graph with accessible interfaces, supporting reproducible knowledge discovery and downstream modelling.

## Methods

### Data sources and libraries

CROssBARv2 is developed to integrate biomedical datasets from multiple sources into a single, unified resource that establishes meaningful relationships across data types to support a wide range of biological data analyses (**Figure 1a**). The system is designed to automatically retrieve and integrate data from relevant sources at regular intervals, ensuring users have continuous access to the latest information. Table S3 and S4 provide a detailed summary of the data sources for each biological entity and relationship integrated into CROssBARv2. Please refer to the **Supplementary Information S8** for detailed information on data sources used in CROssBARv2.

A core component of CROssBARv2’s data infrastructure is Pypath, a Python module designed to process biological data resources by integrating them into comprehensive databases (https://github.com/saezlab/pypath). It offers a versatile Python interface and supports data export, enabling access through other platforms. Designed for modular and standardised access to a wide range of biological databases, pypath is the backbone of OmniPath^56,57^, which integrates over 200 resources. Given its broad compatibility, pypath provides an ideal framework for CROssBARv2’s automated data retrieval workflow. In collaboration with the pypath development team, we extended the tool to support all data sources essential to CROssBARv2. For databases not initially supported, new modules were developed, while existing ones were updated to ensure they retrieved the most complete and current datasets (https://pypath.omnipathdb.org).

CROssBARv2 also employs BioCypher ^8^, an open-source, Python-based platform that enables the swift assembly and upkeep of biomedicalKGs. It simplifies the integration of diverse biomedical datasets by leveraging a modular pipeline that features data ingestion via customizable adapters, schema configuration with standardised ontologies (e.g., the Biolink model ^65^), and comprehensive provenance tracking (https://github.com/biocypher/biocypher). Detailed information on how these data sources and libraries are utilised is provided in the “Building the knowledge graph” section below.

### Building the Knowledge Graph

The CROssBARv2 Knowledge Graph (KG) is stored in a Neo4j graph database, where nodes represent biological entities and edges represent semantic interactions (relationships) among these entities. To construct CROssBARv2 KG, multiple configurable and reusable “adapter” scripts were developed to collect, process, and harmonise biological data (https://github.com/HUBioDataLab/CROssBARv2-KG) (**Figure 1b**). These adapters employ the Pypath library (https://github.com/saezlab/pypath) for data retrieval. Once data is collected, they perform preprocessing and harmonisation to ensure consistency. The processed data is then formatted into node and edge structures compatible with BioCypher, for KG creation. Lastly, BioCypher generated CSV files that were imported into the Neo4j graph database using the Neo4j admin import tool (https://neo4j.com/docs/operations-manual/current/tutorial/neo4j-admin-import/).

For each node type in the KG, common identifiers were utilised to ensure consistency and standardisation. Specifically, we store each node’s identifier in CURIE (compact resource identifier) format (e.g., uniprot:Q9H161) using the prefixes provided by the Bioregistry—an open-source, community-curated meta-registry and identifier resolver ^66^. When an alternative identifier for a biological entity appears in the source data, it is converted to the knowledge graph’s standardised identifier—provided a corresponding cross-reference exists in our data sources. If no such cross-reference is found, the associated data are excluded from the KG. For example, an Ensembl gene identifier is mapped to its corresponding Entrez gene identifier, which serves as the standard for genes in the KG.

In addition to the core entity and relationship data, rich metadata were incorporated as node and edge properties. The complete schema detailing these properties is publicly accessible at https://github.com/HUBioDataLab/CROssBARv2-KG/blob/main/config/schema_config.yaml.

When edges originate from multiple data sources, a deduplication process consolidates each source’s unique properties into a single edge representation while adding a “source” property to record all contributing provenance. For example, if a protein-protein interaction is reported in both the IntAct and STRING databases, the confidence scores from each are retained as separate properties in the unified edge, and both “IntAct” and “STRING” are recorded in its “source” property.

High-quality vector representations of nodes in the KG were generated using various deep learning models tailored to produce embeddings for specific biological entities. Embeddings are dense numerical representations that capture semantic and structural characteristics of entities in a low-dimensional space, enabling efficient similarity comparisons and relationship predictions. For each model, appropriate inputs (e.g., amino acid sequences, SELFIES notations) were prepared and embeddings were generated by feeding these inputs into the corresponding models.

Below, we provided detailed information on each node type, its relationships, and the corresponding embedding method(s) utilised in CROssBARv2. The respective data source is also given for each node and edge type.

#### Organism

Organism nodes were retrieved from UniProtKB ^67^ and identified using NCBI Taxonomy ID ^68^. Organism nodes have two types of edges in the KG: Pathogen-disease relationships, obtained from PathoPhenoDB ^69^, and protein-organism links from UniProtKB.

#### Gene

Gene nodes were obtained from UniProtKB and identified using Entrez Gene identifiers. Gene-protein relationships were sourced from UniProtKB. Transcription factor-gene regulatory interactions were integrated from TRRUST ^70^, CollecTRI ^71^, and DoRothEA ^72^. To ensure consistency, relationships with conflicting regulatory modes (i.e., activation, repression, and unknown or activation and repression) were excluded from the KG. In cases where unknown regulation co-occurred with activation or repression, the latter modes were prioritised to maintain unambiguous regulatory annotations. Orthology relationships linking orthologous genes were incorporated from OMA ^73^ and Pharos ^74^. Orthologous relationships were established between human genes and those from model organisms and evolutionarily relevant species, as specified in the orthology adapter script (https://github.com/HUBioDataLab/CROssBARv2-KG/blob/main/bccb/orthology_adapter.py). Drug-gene relationships were extracted from the Comparative Toxicogenomics Database (CTD) ^75^. Gene-disease associations were collected from Open Targets ^76^, KEGG ^77^, DisGeNET ^78^, humsavar (https://ftp.uniprot.org/pub/databases/uniprot/current_release/knowledgebase/variants/humsavar.txt), DISEASES ^79^, and ClinVar ^80^.

Gene embeddings were derived using the Nucleotide Transformer, specifically the 2.5B parameter multi-species model, which was pre-trained on genomes from 850 evolutionarily diverse species ^81^. This model encodes sequence-specific and regulatory context, enabling biologically informative embeddings, making it well-suited for capturing gene-level features relevant across species.

#### Protein

Protein nodes were obtained from UniProtKB and identified using UniProt identifiers. Associations between proteins and organisms were gathered from UniProtKB, proteins and domains from InterPro ^82^, and proteins and Enzyme Commission (EC) numbers from Expasy ^83^, all of which were incorporated into the KG as edges. Protein-protein interactions were incorporated from IntAct ^84^, BioGRID ^85^, and STRING ^86^ databases. Only high-confidence interactions (score >0.7) were included from STRING. Protein-Gene Ontology (GO) root term (i.e., Molecular Function, Cellular Component, and Biological Process) links were extracted from the Gene Ontology Annotation (GOA) database ^87^, with electronic annotations (i.e., IEA) filtered out. Protein-pathway associations were included from Reactome ^88^ and KEGG databases, with electronic annotations from Reactome filtered out. Protein-phenotype relationships were extracted from the Human Phenotype Ontology (HPO) database ^89^. Compound-protein interactions were incorporated from ChEMBL ^90^ and STITCH ^91^. Drug-protein associations were retrieved from DrugBank ^92^, ChEMBL, Pharos, DGIdb ^93^, STITCH, and KEGG data sources.

ProtT5 embeddings ^21^ were downloaded from https://www.UniProt.org/help/downloads#embeddings resource by selecting the reviewed (Swiss-Prot) per-protein level dataset. ESM-2 protein embeddings ^20^ were generated by averaging residue embeddings obtained from Swiss-Prot protein sequences with the pretrained “esm2_t36_3B_UR50D” model.

#### Protein Domain

Protein domain nodes were retrieved from the InterPro database and identified using InterPro identifiers. Protein domain-GO root term mappings were integrated from the InterPro2GO dataset. Protein-domain associations were incorporated from InterPro. Domain node embeddings were generated using dom2vec method ^94^, which applies the word2vec algorithm ^95^ by modelling protein sequences as sentences and treating domains as words, resulting in 50-dimensional feature representations.

#### Gene Ontology Terms

Gene Ontology (GO) terms, that is, Molecular Function, Cellular Component, and Biological Process, were obtained from the GOA database and identified using GO accessions. Hierarchical relationships between these terms were also extracted from the GOA database. Protein-GO root term links were extracted from the GOA database, with electronic annotations (i.e., IEA) filtered out.

GO term embeddings were generated using the anc2vec method ^96^, which captures the structural properties of GO terms, including ontological uniqueness, ancestral relationships, and sub-ontology classification, resulting in 200-dimensional vector representations.

#### EC number

Enzyme Commission (EC) numbers were retrieved from Expasy and identified using EC codes. Protein-EC number relationships were also sourced from Expasy. Hierarchical relationships between EC numbers were established by connecting EC classes to their respective subclasses following the standard numerical notation system (e.g., EC 1.1.-.- is a subclass of EC 1.-.-.-).

EC number nodes were encoded as 256-dimensional embeddings derived from the SMILES representations ^97^ of their associated reactions, computed using the Python RXNFP library^98^.

#### Pathway

Metabolic, signaling, and disease pathways were obtained from Reactome and KEGG using corresponding database identifiers. Protein-pathway associations were included from Reactome and KEGG databases, with electronic annotations from Reactome filtered out. Drug-pathway associations were incorporated from both Reactome and KEGG databases, while disease-pathway associations were integrated from KEGG and CTD. Hierarchical relationships between pathways were collected from Reactome. Mappings between Reactome and KEGG pathways were retrieved from ComPath ^99^. Pathway orthology relationships between human and other organisms were established by leveraging shared identifier components across species-specific entries in Reactome and KEGG.

Pathway nodes were embedded as 200-dimensional vectors using the TransE knowledge graph embedding method ^100^, trained on gene-pathway interaction networks via the Python BioKEEN library ^101^.

#### Compound

Drug-like small molecule compounds were retrieved from ChEMBL and identified using ChEMBL identifiers. Compound-protein interactions were incorporated from ChEMBL and STITCH. For compound-protein pairs with multiple bioactivity measurements, median values were calculated for relevant parameters (e.g., IC50, pChEMBL) to ensure a consistent and robust representation of interaction strength.

768-dimensional embeddings for compound nodes were generated from their SELFIES representations using the SELFormer chemical language model ^22^.

#### Drug

Drugs were collected from DrugBank and identified using corresponding identifiers. Drug-protein associations were retrieved from DrugBank, ChEMBL, Pharos, DGIdb, STITCH, and KEGG data sources. For drug-protein pairs with multiple bioactivity measurements, median values were calculated for relevant parameters (e.g., IC50, pChEMBL). Drug-drug interactions were obtained from DDInter and KEGG. Drug-gene relationships were extracted from CTD. Cases of contradictory expression annotations, where a drug was reported to both increase and decrease expression of the same gene, were excluded from the KG to maintain data consistency. Drug-pathway associations were incorporated from both Reactome and KEGG. Drug-side effect associations were incorporated from SIDER ^102^, OffSIDES ^103^, and ADReCS ^104^. Drug indications were retrieved from ChEMBL, CTD, and KEGG.

768-dimensional embeddings for drug nodes were generated from their SELFIES representations using the SELFormer chemical language model ^22^.

#### Side Effect

Side effect nodes were obtained from SIDER, OffSIDES, and ADReCS and identified using MedDRA codes. Drug-side effect associations were incorporated from the aforementioned 3 sources. Hierarchical relationships between side effects were derived from the adverse drug reaction hierarchy provided by the ADReCS.

#### Disease

Diseases were retrieved from Mondo Disease Ontology (Mondo) ^105^ and identified using corresponding identifiers. Gene-disease associations were collected from Open Targets, KEGG, DisGeNET, humsavar, DISEASES, and ClinVar. Disease-disease associations were sourced from DisGeNET, hierarchical relationships were derived from Mondo, and comorbidity interactions were obtained from MalaCards ^106^. Drug indications were retrieved from ChEMBL, CTD, and KEGG. Pathogen-disease relationships were obtained from PathoPhenoDB. Disease-pathway associations were integrated from KEGG and CTD. Phenotype-disease relationships were derived from the HPO.

Disease nodes were represented as 100-dimensional embeddings using the doc2vec method ^107^, generated from their textual descriptions sourced from the PrimeKG dataset ^4^. When descriptions were unavailable in PrimeKG, they were retrieved from the original databases where the nodes were obtained. If no description was available, disease names were used as input.

#### Phenotype

Phenotypes were retrieved from the Human Phenotype Ontology (HPO) and identified using corresponding identifiers. Protein-phenotype, phenotype-disease, and phenotype-phenotype hierarchical relationships were derived from the HPO, with electronically inferred phenotype-disease associations removed.

The node2vec algorithm ^108^, was applied to gene-phenotype relationship networks using the CADA tool ^109^ to generate 160-dimensional embeddings for phenotype nodes.

### Deployment of CROssBARv2

#### Server Infrastructure

A Virtual Private Server (VPS) equipped with Docker (https://www.docker.com/) containerisation is employed to deploy the CROssBARv2 system. The landing page, Neo4j (https://neo4j.com/) Console, GraphQL API (https://graphql.org/) interface, and the Large Language Model (LLM) tool are served by individual Docker containers orchestrated through a combined Docker Compose configuration.

The implementation of containerisation enabled rapid recovery of services in the event of failure, while our adoption of a microservices architecture further ensured system robustness. By isolating individual processes within each service, we were able to decouple them, thereby ensuring that the failure of one component would not impact the operation of others.

Traefik (https://github.com/traefik/traefik) is utilised as a reverse proxy, providing automated SSL certificate renewal, routing, and load-balancing capabilities. Furthermore, GitHub Actions are leveraged to automate the build of all individual Docker images, thereby establishing an efficient Continuous Integration/Continuous Deployment (CI/CD) pipeline.

#### Neo4j Browser

The official Neo4j Docker image (https://hub.docker.com/_/neo4j/) from Docker Hub provides an easy-to-access platform for researchers to interact with CROssBARv2 (**Figure 2**). To enable seamless integration with our LLM tool, we have enabled the APOC plugin within this containerised environment. Furthermore, due to security considerations, we have configured the database in read-only mode to prevent unintended modifications.

#### GraphQL API

To facilitate programmatic access to our graph database, we set up an Application Programming Interface (API). We opted for GraphQL due to its inherent advantages in several key aspects (**Figure 2**). GraphQL enables users to specify the exact attributes they require, including nested relationships and features. For instance, a user can choose to fetch only the “mass” property of a protein or omit it altogether. Moreover, Neo4j provides official support for integrating GraphQL capabilities into their graph databases, which aligns with our implementation goals. By adopting GraphQL, we leverage its design philosophy as a flexible and powerful query language that caters to diverse user needs. In our system, Apollo (https://www.apollographql.com/) serves as the client-facing interface that interacts with our GraphQL API.

#### CROssBAR-LLM

The LLM tool provides a platform for querying and analysing CROssBARv2 KG (**Figure 2**), leveraging a hybrid architecture of a Python-based backend and a React.js frontend. The backend, built using the FastAPI framework (https://fastapi.tiangolo.com/), serves as the core computational engine, orchestrating interactions between the LLMs via Langchain (https://www.langchain.com/) and the Neo4j graph database.

The conversion of natural language questions into executable Cypher queries, the execution of these queries against the Neo4j database and the structuring of the returned results into a human-readable format, is handled entirely within the backend system. The LangChain (https://www.langchain.com/) framework is employed within the backend specifically for managing LLM-related tasks, including prompt handling, model interaction, and integration into the overall query execution workflow. The system prompts used for LLM interactions are defined in the https://github.com/HUBioDataLab/CROssBAR_LLM/blob/main/crossbar_llm/backend/tools/qa_templates.py. The backend also features methods for dynamically generating and validating CSRF tokens. This allows the backend to protect against CSRF attacks, which is essential for maintaining security and data integrity.

The frontend, built with React.js (https://react.dev/) and utilising Material UI (https://mui.com/material-ui/) for component design, presents an intuitive interface. The auto-completion feature is powered by the Fuse.js (https://www.fusejs.io/) utility module, which utilises entity names extracted from the KG. Users can choose from multiple LLM providers, set their respective API keys, and configure various parameters such as the top-K results to return. Vector similarity search is facilitated by the VectorSearch component, which includes the VectorUpload component, allowing users to specify vector categories, upload embeddings, and adjust other parameters. The interface also includes a dedicated ResultsDisplay component, which structures the output from the backend for clear interpretation. This output includes both the generated Cypher query (rendered using react-syntax-highlighter for better readability), the raw query results in JSON format, and a natural language response derived from the LLM. The LatestQueries component stores the most recently executed queries, allowing users to reuse past queries and view their specific parameters.

### Exploration and Benchmarking of CROssBARv2

#### Elucidating Potential Therapeutic Mechanisms of De Novo Molecules via CROssBARv2’s Hybrid Approach

De novo molecules were designed based on MRI-1569, a dual-target drug candidate for cannabinoid receptor 1 and inducible nitric oxide synthase proteins ^35^, as a template, with structural modifications introduced through atom-level alterations.

#### The generation of metapath2vec embeddings

We conducted several use-case analyses to assess whether the diverse biological data sources were cohesively integrated within the KG. To carry out this evaluation, we utilised metapath2vec ^23^, an algorithm specifically designed to learn node embeddings within heterogeneous graphs, where multiple types of nodes and relationships coexist. Metapath2vec captures the complexity of multi-type relationships by using predefined meta-paths, which define sequences of meaningful connections between different types of nodes. By conducting random walks guided by these meta-paths, the algorithm constructs heterogeneous neighbourhoods around each node, capturing both direct and contextually relevant indirect relationships. These neighbourhoods are then encoded into continuous vector representations using a skip-gram model, which maximizes the probability of a node’s context based on its neighbours. We utilised the metapath2vec implementation in PyTorch Geometric ^110^, with key hyperparameters optimised for each use-case—random walk length, number of neighbouring nodes for context, number of random walks per node, count of negative samples, and training batch size—resulting in 128-dimensional node embeddings.

#### LLM Benchmarks

For the Cypher query generation task, a benchmark set of 50 questions along with their corresponding ground truth Cypher queries was manually created by a Neo4j expert. Since some questions could return an extremely high number of results, all Cypher queries generated by LLMs were constrained to return a maximum of five outputs (i.e., data points). A generated query was considered correct if its results constituted a complete subset of the ground truth answers.

A true-false evaluation set comprising 100 questions was curated from gene-disease and drug-disease relationships within the KG. For the negative set, node pairs were randomly selected from the same categories and included only if no direct relationship between them was found. The user prompt provided to the LLMs for these questions was: “{question}. Return only tf_effect.”

GeneTuring benchmark was downloaded from https://raw.githubusercontent.com/monarch-initiative/oai-plugin-evals/main/datasets/geneturing/geneturing_converted.csv. The user prompt provided to the LLMs for these questions was: “{question}. Return only tf_effect.” For LLaMA 3.1-405B, a slightly modified prompt was used: “{question}. Return only tf_effect property of relationship.”

The responses generated by the LLMs for all tasks, along with the questions and ground truth answers for the first two tasks, are provided as **Supplementary Files**.

### Performance Metrics

To evaluate the quality of the generated node embeddings and their ability to capture biologically meaningful relationships, we employed multiple performance metrics. We calculated cosine similarity between nodes to quantitatively assess the strength of relationships captured in the embeddings. This score indicates how closely related two nodes are, with higher values reflecting stronger relationships and lower values suggesting weaker or no connections.

Building upon cosine similarity, we assessed ranking accuracy using the Hit Rate at K (HR@K) metric. For each edge type in our use-cases, we generated a ranked list of candidate edges based on a scoring function (e.g., cosine similarity between the source and target node embeddings). HR@K measures the proportion of ground-truth relations that are recovered within the top-K ranked predictions, normalized by the maximum number of true relations that could possibly appear in the top-K (Eq. 1).

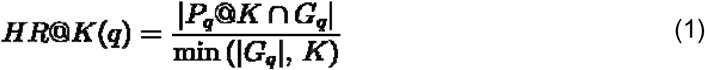

where, *G_q_* is the set of ground-truth relations for the edge type *q*; and *P_q_@K* is the set of top-K predicted relationships. The overall HR@K score for an edge type is obtained by averaging HR@K across all queries. Higher HR@K values indicate better alignment between embedding-based similarity and the true relational structure of the knowledge graph.

To further evaluate the embeddings, we performed clustering analysis to examine how well biologically similar entities grouped together. In the protein relationships use-case, we focused on whether proteins from the same family formed coherent groups in the embedding space. To stabilize the analysis, embeddings were first reduced to 50 dimensions using PCA and then L2-normalized to unit length. We then applied agglomerative hierarchical clustering with average linkage and cosine distance, using the implementation from the scikit-learn library ^111^, and set the number of clusters equal to the number of protein families considered. Clustering quality was quantified using purity, defined as the proportion of proteins correctly assigned to the dominant family label within each cluster. To visualize the clustering patterns, we used t-SNE ^112^, which projects high-dimensional data into two dimensions for easier interpretation, employing the scikit-learn implementation.

We next examined CROssBARv2’s utility for downstream predictive tasks, by training deep learning models to predict protein functions. For evaluating prediction performance, we used the widely adopted Fmax metric. Fmax is calculated from probabilistic outputs generated by the model across varying decision thresholds, where predictions above a given threshold are treated as positive and those below as negative. It represents the maximum F1-score (Eq. 2) achieved across all thresholds, where F1 is the harmonic mean of precision (pr) and recall (rc).

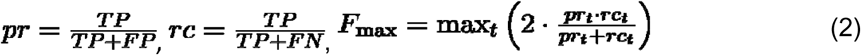

*TP*, *FP*, *FN*, and *TN* correspond to true positives, false positives, false negatives, and true negatives, respectively. The threshold level is denoted by *t*.

## Data availability

The CROssBARv2 project website is available at https://crossbarv2.hubiodatalab.com/, providing access to the GraphQL API, Neo4j Browser, and CROssBAR-LLM tool (please use: https://crossbarv2.hubiodatalab.com/llm). Neo4j-importable CSV files required to reconstruct the KG are publicly available at https://drive.google.com/file/d/1KoMAxlvy_4IOo8MPi4TrSbMlQtBf8Pch/view?usp=sharing. The node embeddings stored in the graph database and used for semantic similarity search are also publicly available at https://drive.google.com/file/d/1HRUlQ_PaunSH7Rs8ZCihKdiVxiOZrE0W/view?usp=sharing. In addition, the raw node and edge datasets underlying the CROssBARv2 KG are hosted on Hugging Face at https://huggingface.co/datasets/HUBioDataLab/CROssBARv2-KG.

## Code availability

The source code for KG construction adapters, GraphQL API, CROssBAR-LLM, and Docker Compose deployment scripts is openly available at https://github.com/HUBioDataLab/CROssBARv2 organized as subrepositories.

## Acknowledgements

This project is funded by TUBITAK-ARDEB 3501 Career Development Program under project no. 120E531 and by the European Union’s Horizon 2020 research and innovation programme (grant agreement No 965193, DECIDER). BŞ and EU acknowledge support from the TUBITAK-BIDEB 2210 National MSc/MA Scholarship Program. ASR is supported by the Heidelberg Faculty of Medicine at Heidelberg University through the Medical Data Scientist Fellowship. DT was supported by the Landesinstitut für Bioinformatikinfrastruktur in Baden-Württemberg.

## Author Contributions Statement

TD conceptualized and supervised the project with support from JSR. DT, EU, and MD developed pypath/OmniPath scripts for data retrieval. BŞ and EU prepared adapter scripts for data processing and harmonization. SL and BŞ integrated and implemented BioCypher within the adapter scripts. BŞ built the Neo4j graph database. MD constructed the GraphQL API, implemented the server infrastructure and the project website. ME and BŞ developed the infrastructure for CROssBAR-LLM, and ME implemented its front-end. BŞ and EU conducted use-case analyses, LLM-centric analyses, visualised the results and prepared figures of the article. BŞ, EU, MD, ME, and TD wrote the paper. SL, ASR, DT, JSR, and TD reviewed the article. All authors approved the text.

## References

1. Doğan, T. et al. CROssBAR: comprehensive resource of biomedical relations with knowledge graph representations. Nucleic Acids Res. 49, e96 (2021).

2. Himmelstein, D. S. et al. Systematic integration of biomedical knowledge prioritizes drugs for repurposing. eLife 6, (2017).

3. Morris, J. H. et al. The scalable precision medicine open knowledge engine (SPOKE): a massive knowledge graph of biomedical information. Bioinformatics 39, btad080 (2023).

4. Chandak, P., Huang, K. & Zitnik, M. Building a knowledge graph to enable precision medicine. Scientific Data 10, 1–16 (2023).

5. Fernández-Torras, A., Duran-Frigola, M., Bertoni, M., Locatelli, M. & Aloy, P. Integrating and formatting biomedical data as pre-calculated knowledge graph embeddings in the Bioteque. Nat Commun 13, 5304 (2022).

6. Santos, A. et al. A knowledge graph to interpret clinical proteomics data. Nat. Biotechnol. 40, 692–692 (2022).

7. DoLan, T. et al. CROssBAR: comprehensive resource of biomedical relations with knowledge graph representations. Nucleic Acids Res. 49, e96–e96 (2021).

8. Lobentanzer, S. et al. Democratizing knowledge representation with BioCypher. Nat. Biotechnol. 41, 1056–1059 (2023).

9. Zheng, S. et al. PharmKG: a dedicated knowledge graph benchmark for bomedical data mining. Brief. Bioinform. 22, (2021).

10. Gema, A. P., et al. Knowledge graph embeddings in the biomedical domain: Are they useful? A look at link prediction, rule learning, and downstream polypharmacy tasks. arXiv [cs.LG] (2023) doi:10.48550/ARXIV.2305.19979.

11. Vilela, J. et al. Biomedical knowledge graph embeddings for personalized medicine: Predicting diseaseLgene associations. Expert Syst. 40, (2023).

12. Xu, Z., Jain, S. & Kankanhalli, M. Hallucination is Inevitable: An Innate Limitation of Large Language Models. Preprint at 10.48550/ARXIV.2401.11817 (2024).

13. Perez, E. et al. Discovering language model behaviors with model-written evaluations. arXiv [cs.CL] (2022) doi:10.48550/ARXIV.2212.09251.

14. Introducing deep research. https://openai.com/index/introducing-deep-research/ (2025).

15. Introducing the Model Context Protocol. https://www.anthropic.com/news/model-context-protocol (2024).

16. Miller, H. E., Greenig, M., Tenmann, B. & Wang, B. BioML-bench: Evaluation of AI agents for end-to-end biomedical ML. bioRxiv (2025) doi:10.1101/2025.09.01.673319.

17. Agrawal, G., Kumarage, T., Alghamdi, Z. & Liu, H. Can knowledge graphs reduce hallucinations in LLMs? : A survey. arXiv [cs.CL] (2023) doi:10.48550/ARXIV.2311.07914.

18. Pan, S. et al. Unifying large language models and Knowledge Graphs: A roadmap. arXiv [cs.CL] (2023) doi:10.48550/ARXIV.2306.08302.

19. Neo4j Graph Database. Graph Database & Analytics https://neo4j.com/ (2020).

20. Lin, Z. et al. Evolutionary-scale prediction of atomic-level protein structure with a language model. Science 379, 1123–1130 (2023).

21. Elnaggar, A., et al. ProtTrans: Towards cracking the language of life’s code through self-supervised deep learning and high performance computing. arXiv [cs.LG] (2020) doi:10.48550/ARXIV.2007.06225.

22. Yüksel, A., Ulusoy, E., Ünlü, A. & Doğan, T. SELFormer: molecular representation learning via SELFIES language models. Mach. Learn.: Sci. Technol. 4, 025035 (2023).

23. Dong, Y., Chawla, N. V. & Swami, A. metapath2vec: Scalable Representation Learning for Heterogeneous Networks. in Proceedings of the 23rd ACM SIGKDD International Conference on Knowledge Discovery and Data Mining 135–144 (Association for Computing Machinery, New York, NY, USA, 2017).

24. Atas Guvenilir, H. & Doğan, T. How to approach machine learning-based prediction of drug/compound–target interactions. Journal of Cheminformatics 15, 16 (2023).

25. Schwartz, R. S. et al. Human recombinant DNA-derived antihemophilic factor (factor VIII) in the treatment of hemophilia A. recombinant Factor VIII Study Group. N. Engl. J. Med. 323, 1800–1805 (1990).

26. Lusher, J. et al. Human recombinant DNA-derived antihemophilic factor in the treatment of previously untreated patients with hemophilia A: final report on a hallmark clinical investigation. Journal of thrombosis and haemostasis : JTH 2, (2004).

27. Engeli, S. et al. Activation of the peripheral endocannabinoid system in human obesity. Diabetes 54, 2838–2843 (2005).

28. Ghasemi-Gojani, E., Kovalchuk, I. & Kovalchuk, O. Cannabinoids and terpenes for diabetes mellitus and its complications: from mechanisms to new therapies. Trends Endocrinol. Metab. 33, 828–849 (2022).

29. Barutta, F., Mastrocola, R., Bellini, S., Bruno, G. & Gruden, G. Cannabinoid receptors in diabetic kidney disease. Curr. Diab. Rep. 18, 9 (2018).

30. Mukhopadhyay, P. et al. CB1 cannabinoid receptors promote oxidative stress and cell death in murine models of doxorubicin-induced cardiomyopathy and in human cardiomyocytes. Cardiovasc. Res. 85, 773–784 (2010).

31. Rajesh, M. et al. Cannabinoid 1 receptor promotes cardiac dysfunction, oxidative stress, inflammation, and fibrosis in diabetic cardiomyopathy. Diabetes 61, 716–727 (2012).

32. Noronha, B. T., Li, J.-M., Wheatcroft, S. B., Shah, A. M. & Kearney, M. T. Inducible nitric oxide synthase has divergent effects on vascular and metabolic function in obesity. Diabetes 54, 1082–1089 (2005).

33. Król, M. & Kepinska, M. Human nitric oxide synthase-its functions, polymorphisms, and inhibitors in the context of inflammation, diabetes and cardiovascular diseases. Int. J. Mol. Sci. 22, 56 (2020).

34. Mungrue, I. N. et al. Cardiomyocyte overexpression of iNOS in mice results in peroxynitrite generation, heart block, and sudden death. J. Clin. Invest. 109, 735–743 (2002).

35. Cinar, R., Iyer, M. R. & Kunos, G. The therapeutic potential of second and third generation CB1R antagonists. Pharmacol. Ther. 208, 107477 (2020).

36. Roger, C. et al. Simultaneous inhibition of peripheral CB1R and iNOS mitigates obesity-related dyslipidemia through distinct mechanisms. Diabetes 69, 2120–2132 (2020).

37. Hou, W., Shang, X. & Ji, Z. Benchmarking large language models for genomic knowledge with GeneTuring. bioRxivorg (2025) doi:10.1101/2023.03.11.532238.

38. Sonar by Perplexity. https://sonar.perplexity.ai/.

39. Ulusoy, E. & Doğan, T. ProtHGT: Heterogeneous graph transformers for automated protein function prediction using biological knowledge graphs and language models. bioRxiv (2025) doi:10.1101/2025.04.19.649272.

40. Zhao, Y. et al. Predicting Protein Functions Based on Heterogeneous Graph Attention Technique. IEEE J Biomed Health Inform 28, 2408–2415 (2024).

41. Soroush, A., et al. Large language models are poor medical coders — benchmarking of medical code querying. NEJM AI 1, (2024).

42. Doneva, S. E. et al. Large language models to process, analyze, and synthesize biomedical texts: a scoping review. *Discov*. Artif. Intell. 4, (2024).

43. Kuehl, M. et al. BioContextAI is a community hub for agentic biomedical systems. Nat Biotechnol 43, 1755–1757 (2025).

44. Gao, S. et al. Empowering biomedical discovery with AI agents. Cell 187, 6125–6151 (2024).

45. Papatheodorou, I. et al. Expression Atlas update: from tissues to single cells. Nucleic Acids Res. 48, D77–D83 (2020).

46. GTEx Consortium. The GTEx Consortium atlas of genetic regulatory effects across human tissues. Science 369, 1318–1330 (2020).

47. Huang, Q., Szklarczyk, D., Wang, M., Simonovic, M. & von Mering, C. PaxDb 5.0: Curated protein quantification data suggests adaptive proteome changes in yeasts. Mol. Cell. Proteomics 22, 100640 (2023).

48. Lautenbacher, L. et al. ProteomicsDB: toward a FAIR open-source resource for life-science research. Nucleic Acids Res. 50, D1541–D1552 (2022).

49. Metabolomics Workbench. https://www.metabolomicsworkbench.org/.

50. Wishart, D. S. et al. HMDB 5.0: The Human Metabolome Database for 2022. Nucleic Acids Res. 50, D622–D631 (2022).

51. ENCODE Project Consortium. The ENCODE (ENCyclopedia Of DNA elements) Project. Science 306, 636–640 (2004).

52. Bernstein, B. E. et al. The NIH Roadmap Epigenomics Mapping Consortium. Nat. Biotechnol. 28, 1045–1048 (2010).

53. Zhang, M. et al. MethBank 4.0: an updated database of DNA methylation across a variety of species. Nucleic Acids Res. 51, D208–D216 (2023).

54. CZI Cell Science Program et al. CZ CELLxGENE Discover: a single-cell data platform for scalable exploration, analysis and modeling of aggregated data. Nucleic Acids Res. 53, D886–D900 (2025).

55. Richardson, L. et al. MGnify: the microbiome sequence data analysis resource in 2023. Nucleic Acids Res. 51, D753–D759 (2023).

56. Türei, D., Korcsmáros, T. & Saez-Rodriguez, J. OmniPath: guidelines and gateway for literature-curated signaling pathway resources. Nat Methods 13, 966–967 (2016).

57. Türei, D. et al. OmniPath: integrated knowledgebase for multi-omics analysis. Nucleic Acids Res (2025) doi:10.1093/nar/gkaf1126.

58. Scalbert, A. et al. Databases on food phytochemicals and their health-promoting effects. J. Agric. Food Chem. 59, 4331–4348 (2011).

59. Dooley, D. M. et al. FoodOn: a harmonized food ontology to increase global food traceability, quality control and data integration. Npj Sci. Food 2, 23 (2018).

60. Hayes, T. et al. Simulating 500 million years of evolution with a language model. Science 387, 850–858 (2025).

61. Brixi, G. et al. Genome modeling and design across all domains of life with Evo 2. bioRxiv (2025) doi:10.1101/2025.02.18.638918.

62. Zhou, G. et al. Uni-Mol: A universal 3D molecular representation learning framework. ChemRxiv (2023) doi:10.26434/chemrxiv-2022-jjm0j-v4.

63. Ji, X., et al. Uni-Mol2: Exploring molecular pretraining model at scale. arXiv [cs.LG] (2024) doi:10.48550/ARXIV.2406.14969.

64. Cai, Y., Guo, Z., Pei, Y., Bian, W. & Zheng, W. SimGRAG: Leveraging similar subgraphs for knowledge graphs driven retrieval-Augmented Generation. arXiv [cs.CL] (2024) doi:10.48550/ARXIV.2412.15272.

65. Unni, D. R. et al. Biolink Model: A universal schema for knowledge graphs in clinical, biomedical, and translational science. arXiv [cs.DB] (2022) doi:10.48550/ARXIV.2203.13906.

66. Hoyt, C. T. et al. Unifying the identification of biomedical entities with the Bioregistry. Sci. Data 9, 714 (2022).

67. UniProt Consortium. UniProt: The universal protein knowledgebase in 2023. Nucleic Acids Res. 51, D523–D531 (2023).

68. Schoch, C. L. et al. NCBI Taxonomy: a comprehensive update on curation, resources and tools. Database (Oxford) 2020, (2020).

69. Kafkas, Ş., et al. PathoPhenoDB, linking human pathogens to their phenotypes in support of infectious disease research. Sci. Data 6, 79 (2019).

70. Han, H. et al. TRRUST v2: an expanded reference database of human and mouse transcriptional regulatory interactions. Nucleic Acids Res. 46, D380–D386 (2018).

71. Müller-Dott, S. et al. Expanding the coverage of regulons from high-confidence prior knowledge for accurate estimation of transcription factor activities. Nucleic Acids Res. 51, 10934–10949 (2023).

72. Garcia-Alonso, L., Holland, C. H., Ibrahim, M. M., Turei, D. & Saez-Rodriguez, J. Benchmark and integration of resources for the estimation of human transcription factor activities. Genome Res. 29, 1363–1375 (2019).

73. Altenhoff, A. M. et al. OMA orthology in 2024: improved prokaryote coverage, ancestral and extant GO enrichment, a revamped synteny viewer and more in the OMA Ecosystem. Nucleic Acids Res. 52, D513–D521 (2024).

74. Sheils, T. K. et al. TCRD and Pharos 2021: mining the human proteome for disease biology. Nucleic Acids Res. 49, D1334–D1346 (2021).

75. Davis, A. P. et al. Comparative Toxicogenomics Database’s 20th anniversary: update 2025. Nucleic Acids Res. 53, D1328–D1334 (2025).

76. Koscielny, G. et al. Open Targets: a platform for therapeutic target identification and validation. Nucleic Acids Res. 45, D985–D994 (2017).

77. Kanehisa, M., Furumichi, M., Tanabe, M., Sato, Y. & Morishima, K. KEGG: new perspectives on genomes, pathways, diseases and drugs. Nucleic Acids Res. 45, D353–D361 (2017).

78. Piñero, J. et al. The DisGeNET knowledge platform for disease genomics: 2019 update. Nucleic Acids Res. 48, D845–D855 (2020).

79. Grissa, D., Junge, A., Oprea, T. I. & Jensen, L. J. Diseases 2.0: a weekly updated database of disease-gene associations from text mining and data integration. Database (Oxford) 2022, (2022).

80. Landrum, M. J. et al. ClinVar: public archive of relationships among sequence variation and human phenotype. Nucleic Acids Res. 42, D980–5 (2014).

81. Dalla-Torre, H., et al. The Nucleotide Transformer: Building and evaluating robust foundation models for human genomics. bioRxiv (2023) doi:10.1101/2023.01.11.523679.

82. Blum, M. et al. InterPro: the protein sequence classification resource in 2025. Nucleic Acids Res. 53, D444–D456 (2025).

83. Duvaud, S. et al. Expasy, the Swiss Bioinformatics Resource Portal, as designed by its users. Nucleic Acids Res. 49, W216–W227 (2021).

84. Del Toro, N. et al. The IntAct database: efficient access to fine-grained molecular interaction data. Nucleic Acids Res. 50, D648–D653 (2022).

85. Oughtred, R. et al. The BioGRID database: A comprehensive biomedical resource of curated protein, genetic, and chemical interactions. Protein Sci. 30, 187–200 (2021).

86. Szklarczyk, D. et al. The STRING database in 2023: protein-protein association networks and functional enrichment analyses for any sequenced genome of interest. Nucleic Acids Res. 51, D638–D646 (2023).

87. Gene Ontology Consortium et al. The Gene Ontology knowledgebase in 2023. Genetics 224, (2023).

88. Milacic, M. et al. The reactome pathway knowledgebase 2024. Nucleic Acids Res. 52, D672–D678 (2024).

89. Köhler, S. et al. The Human Phenotype Ontology in 2021. Nucleic Acids Res. 49, D1207–D1217 (2021).

90. Zdrazil, B. et al. The ChEMBL Database in 2023: a drug discovery platform spanning multiple bioactivity data types and time periods. Nucleic Acids Res. 52, D1180–D1192 (2024).

91. Szklarczyk, D. et al. STITCH 5: augmenting protein-chemical interaction networks with tissue and affinity data. Nucleic Acids Res. 44, D380–4 (2016).

92. Knox, C. et al. DrugBank 6.0: The DrugBank knowledgebase for 2024. Nucleic Acids Res. 52, D1265–D1275 (2024).

93. Cannon, M. et al. DGIdb 5.0: rebuilding the drug-gene interaction database for precision medicine and drug discovery platforms. Nucleic Acids Res. 52, D1227–D1235 (2024).

94. Melidis, D. P. & Nejdl, W. Capturing Protein Domain Structure and Function Using Self-Supervision on Domain Architectures. Algorithms 2021, Vol. 14, Page 28 14, 28–28 (2021).

95. Mikolov, T., Chen, K., Corrado, G. & Dean, J. Efficient Estimation of Word Representations in Vector Space. 1st International Conference on Learning Representations, ICLR 2013 - Workshop Track Proceedings (2013).

96. Edera, A. A., Milone, D. H. & Stegmayer, G. Anc2vec: embedding gene ontology terms by preserving ancestors relationships. Brief. Bioinform. 23, (2022).

97. Weininger, D. SMILES, a Chemical Language and Information System: 1: Introduction to Methodology and Encoding Rules. J. Chem. Inf. Comput. Sci. 28, 31–36 (1988).

98. Schwaller, P. et al. Mapping the space of chemical reactions using attention-based neural networks. Nature Machine Intelligence 2021 3:2 3, 144–152 (2021).

99. Domingo-Fernández, D., Hoyt, C. T., Bobis-Álvarez, C., Marín-Llaó, J. & Hofmann-Apitius, M. ComPath: an ecosystem for exploring, analyzing, and curating mappings across pathway databases. NPJ Syst. Biol. Appl. 5, 3 (2019).

100. Bordes, A., Usunier, N., Garcia-Duran, A., Weston, J. & Yakhnenko, O. Translating Embeddings for Modeling Multi-relational Data. in Advances in Neural Information Processing Systems vol. 26 (Curran Associates, Inc., 2013).

101. Ali, M., Hoyt, C. T., Domingo-Fernández, D., Lehmann, J. & Jabeen, H. BioKEEN: a library for learning and evaluating biological knowledge graph embeddings. Bioinformatics 35, 3538–3540 (2019).

102. Kuhn, M., Letunic, I., Jensen, L. J. & Bork, P. The SIDER database of drugs and side effects. Nucleic Acids Res. 44, D1075–9 (2016).

103. Tatonetti, N. P., Ye, P. P., Daneshjou, R. & Altman, R. B. Data-driven prediction of drug effects and interactions. Sci. Transl. Med. 4, 125ra31 (2012).

104. Cai, M.-C. et al. ADReCS: an ontology database for aiding standardization and hierarchical classification of adverse drug reaction terms. Nucleic Acids Res. 43, D907–13 (2015).

105. Vasilevsky, N. A., et al. Mondo: Unifying diseases for the world, by the world. bioRxiv (2022) doi:10.1101/2022.04.13.22273750.

106. Rappaport, N. et al. MalaCards: an amalgamated human disease compendium with diverse clinical and genetic annotation and structured search. Nucleic Acids Res. 45, D877–D887 (2017).

107. Řeh uřek, R. & Sojka, P. Software Framework for Topic Modelling with Large Corpora. (ELRA, 2010).

108. Grover, A. & Leskovec, J. node2vec: Scalable Feature Learning for Networks. Proceedings of the ACM SIGKDD International Conference on Knowledge Discovery and Data Mining 13-17-August-2016, 855–864 (2016).

109. Peng, C. et al. CADA: phenotype-driven gene prioritization based on a case-enriched knowledge graph. NAR Genom. Bioinform. 3, (2021).

110. Fey, M. & Lenssen, J. E. Fast Graph Representation Learning with PyTorch Geometric. (2019).

111. Pedregosa, F. et al. Scikit-learn: Machine Learning in Python. Journal of Machine Learning Research (2011).

112. Van der Maaten, L. & Hinton, G. Visualizing data using t-SNE. J. Mach. Learn. Res. 9, (2008).

