## Supplementary Information for "CROssBARv2: A Unified Computational Framework for Heterogeneous Biomedical Data Representation and LLM-Driven Exploration"

### S1. Extended Introduction

Advances in biomedical research increasingly rely on data-driven approaches to unravel complex systems and develop solutions for challenging diseases. The ability to analyse large-scale biological and biomedical data is essential for identifying underlying mechanisms and supporting personalised therapeutic strategies. However, biological data remains highly fragmented across numerous specialised resources. Institutions often curate specific datasets in isolation, leaving connections between these datasets implicit or poorly defined. This lack of interconnectivity limits the discovery of meaningful relationships and prevents the full exploitation of the complementary information these diverse datasets offer [1]. Addressing this issue requires robust systems that integrate, organise, and model these datasets into unified frameworks. These frameworks facilitate the analysis of complex biological interactions, thereby empowering researchers to identify insights that remain elusive in isolated datasets. With the systematic modelling of biological entities and their interrelationships, we can gain deeper insights into biological systems and accelerate progress in precision medicine and therapeutic development.

Knowledge graphs (KGs) have emerged as powerful tools for addressing these challenges. By representing biological entities—such as genes, proteins, pathways, diseases, drugs, etc.—as nodes and their interactions as edges, KGs provide a comprehensive, interconnected network. They seamlessly integrate diverse datasets while preserving contextual relationships, effectively capturing the complexity of biological systems. Beyond offering an intuitive visualisation of biological interactions, KGs support advanced computational analyses, including path-based similarity, hypothesis generation, and discovery of hidden relationships. With their flexibility and scalability, KGs have become [2] essential for managing the growing complexity of biological data and supporting innovative research [3].

Several KGs have been developed so far to combine and analyse biomedical data, each addressing specific aspects of data integration and application in biomedical research. Hetionet, a pioneering biomedical KG, systematically includes diverse biomedical datasets to support drug repurposing efforts [4]. By leveraging a richly interconnected structure of the KGs, Hetionet enables predictions of therapeutic connections between drugs and diseases while supporting complex queries across pathways, side effects, and gene functions. Comprising 21 node and 55 edge types, SPOKE spans domains from molecular biology to clinical practice [5] and supports graph-based machine learning and predictive modelling. PrimeKG integrates data from 20 high-quality sources to support precision medicine applications [2]. Covering ten biological entities, including molecular, cellular, tissue, and population levels, it offers a multiscale biological representation. This framework links diseases, drugs, phenotypes, and other entities, focusing on drug-disease relationships like indications and off-label uses. Bioteque accommodates biomedical data from over 150 sources and incorporates pre-calculated node embeddings, summarising complex relationships for machine learning applications [6]. This KG is purported to facilitate tasks such as drug repurposing, pathway identification, and gene-disease association prediction. The Clinical Knowledge Graph (CKG) specialises in clinical proteomics and other omics data [7]. By consolidating biomedical databases, experimental results, and scientific literature, CKG provides a graph-based structure optimised for proteomics workflows. CROssBARv1, the precursor to this study, demonstrated the potential of automated and systematic integration of heterogeneous biomedical data into a KG structure [1]. It offered a user-friendly interface for generating knowledge graphs on the fly based on simple queries, aiding the visualisation of complex biological networks. CROssBARv1's utility was particularly evident in systems-level research and drug discovery, such as exploring protein-protein interactions, gene/protein-pathway relationships, phenotype-assisted disease mechanisms, and drug/compound-target interactions, including deep learning-based predictions. To assist biomedical KG construction, the open-source BioCypher framework further lowers the barrier to entry by providing a FAIR-compliant pipeline that automates ingestion, semantic harmonisation and export of heterogeneous biomedical data into graph back-ends such as Neo4j, complete with provenance tracking [3].

While progress has been made in unifying heterogeneous datasets within integrated platforms, significant limitations remain in functionality, maintainability, reproducibility, and accessibility. Many current KGs lack the flexibility to address specialised research needs and broader applications. For instance, the Drug Repurposing Knowledge Graph (DRKG) [8] is explicitly tailored for drug development initiatives, while PrimeKG is designed to support precision medicine. Such specialised designs, often with limited node and edge types, constrain utility across diverse use cases. Compounding these limitations, existing KGs often lack comprehensive metadata annotations for biological entities, neglecting valuable contextual information such as data provenance, experimental validation status, and confidence scores that could otherwise enhance their reliability, utility, and interpretability and enable more informed decision-making. Usability is another key issue. Many KGs rely on data delivery formats and access methods that inherently restrict usability for those without programming skills. For example, PrimeKG [2] and PharmKG [9] primarily distribute their data as CSV files, targeting programmatic users and requiring coding expertise for data extraction and analysis. Reproducibility and automation are further areas of concern. Numerous influential KGs, including DRKG and PharmKG, do not publicly share their construction code, hindering independent replication and community-driven improvements. Similarly, maintainability remains a critical challenge, as none of the detailed KGs discussed here has a formal maintenance or update schedule in place. A notable example is Hetionet, which has not been updated since 2017. The absence of systematic version tracking and automated update workflows across many KGs further exacerbates these issues, reducing their long-term relevance and adaptability in rapidly evolving scientific domains. When we consider content-related problems, few KGs include critical genomic and transcriptomic layers, such as single-nucleotide polymorphism (SNP) data [10–12], and to our knowledge, none incorporate bulk RNA-seq or cutting-edge single-cell expression datasets. Incorporating such data would significantly enhance the resolution and relevance of KGs, enabling more precise insights into biological systems. Although a few previous studies have explored KG embeddings or random-walk-based methods to learn representations of biological entities directly from KGs [6,13,14], to the best of our knowledge, no existing work has leveraged feature-rich embeddings generated by external representation learning models (e.g., ESM2, ProtTrans, etc.), which could significantly improve query procedures in KGs or any biological database via enabling semantic similarity search. Addressing these gaps is crucial for developing adaptable, user-friendly, and cutting-edge KG-based systems capable of meeting the needs of the biomedical community.

Large language models (LLMs) have emerged as powerful tools for processing and generating natural language, with broad applications across the biological and biomedical fields. Their ability to analyse vast textual datasets has enabled them to extract valuable insights from scientific articles, clinical documentation, and other unstructured sources [15–18]. This makes LLMs contribute significantly to biomedical question answering and retrieving accurate and contextually insightful information. However, LLMs face significant limitations in this domain. Their reliance on pre-trained knowledge from static datasets often results in gaps in domain-specific knowledge and an inability to reflect the latest advancements [19]. They also struggle with factual accuracy [20,21], especially when tasked with resolving highly technical or specialised queries that require detailed contextual understanding [22,23]. Additionally, LLMs lack direct access to structured databases, restricting their ability to validate information or provide precise, data-supported answers. Integrating LLMs with KGs offers a promising solution to these challenges [24–26]. By combining the linguistic capabilities of LLMs with the rich repository of structured, interconnected, and up-to-date data in KGs, this hybrid approach enhances the accuracy, depth, and reliability of question-answering systems [27,28]. KGs can provide LLMs with direct access to verified relationships and detailed attributes of biological entities, allowing for accurate and context-aware responses. This integration may bridge the gap between unstructured text processing and structured data representation, enabling more robust, context-aware, and trustworthy applications in biomedical research.

### S2. Expanded Evaluation of CROssBARv2 Knowledge Graph Use Case Analyses

S2.1. Exploring the functional significance of protein relationships

In the first KG relationship assessment-related use case, we explored the biological significance of activity/role-centric connections of protein nodes. We used metapath2vec to generate embeddings based on protein–pathway, protein–GO term, and protein–EC number relationships. These metapaths were selected to capture fundamental aspects of protein function—participation in biological processes, execution of molecular functions, and catalysis of enzymatic reactions—all of which contribute to a protein's functional identity and are often conserved within protein families. By embedding proteins through these functional properties, we aimed to assess whether proteins from the same families cluster together in the embedding space, reflecting their true functional characteristics as represented by their relationships in the KG. We visualised the embedding space using t-SNE [29], a common technique for dimensionality reduction. Using a dataset of drug target protein superfamilies [30], proteins are grouped into five enzyme classes (hydrolases, oxidoreductases, proteases, transferases, and other enzymes) and five non-enzyme classes (epigenetic regulators, ion channels, membrane receptors, transcription factors, and transporters). The proteins are colour-coded by superfamily in the t-SNE projection to assess how the members of specific families are clustered in the resulting vector space (Figure 2a).

Figure 2a shows that non-enzyme proteins within the same family were clearly clustered, with well-defined separations between families. For enzyme families, the clustering was less distinct, likely due to the broad characteristics shared by the main classes of enzymes and their involvement in a variety of biological processes. To explore these differences in clustering performance, we quantified the shared biological context between proteins by calculating the number of shared edges: cases where two proteins connect to the same pathway, GO term, or EC number. For each protein, we computed the mean number of unique shared edges with proteins from the same family (intra-family) and from different families (inter-family). These values were then averaged across all members of a family to obtain family-level intra- and inter-family means, providing insight into how much functional overlap underlies the observed clustering patterns. Non-enzyme families exhibited approximately 24 times more common edges within the family than with other families (Figure 2b), while enzyme families showed a smaller but still prominent difference of around 8 times.

To establish a comparative baseline, we evaluated CROssBARv2 protein embeddings against Bioteque (Fernández-Torras et al. 2022), a comprehensive repository of pre-computed embeddings derived using metapath-guided node2vec from a KG that integrates more than 150 biomedical data sources. For comparability, we selected the Bioteque embedding type most closely related to our protein representations, namely "GEN-ass-PWY", which encodes gene–pathway associations. Unlike CROssBARv2, Bioteque's network does not include GO term or EC number entity types; consequently, these relationships are not represented in its metapath definitions.

We applied agglomerative hierarchical clustering strategy to both embedding sets, using curated enzyme and non-enzyme protein families as ground-truth labels, and assessed cluster quality with purity, which measures how often proteins from the same family fall into the same cluster. CROssBARv2 achieved purity scores of 0.504 for enzyme families and 0.818 for non-enzymes, compared with 0.445 and 0.773 for Bioteque, respectively.

These findings indicate that CROssBARv2's embedding space reflects biologically meaningful organisation of protein families. By capturing pathways, molecular functions, and enzymatic activities, the KG can reveal hidden patterns and support the functional characterization of unannotated proteins. Comparison with Bioteque further highlights that explicitly integrating GO and EC associations in addition to pathways yields a more complete representation of protein functional organisation.

#### S2.2. Analysing drug-disease associations for therapeutic relevance

In this use case, the overall aim is to explore the potential of our KG for modelling drug and disease relationships by evaluating whether the proximity of drug and disease nodes in the embedding space reflects their true biological associations captured by their diverse relationships in the graph. For this analysis, we selected five edge types: disease-drug associations, drug-drug interactions, and three types of disease-disease edges—hierarchy, association, and comorbidity—to define metapaths for generating embeddings of drug and disease nodes with the metapath2vec model. These edge types were chosen to capture complementary factors relevant to drug applicability and disease context. This was followed by the calculation of pairwise cosine similarities between embeddings of all node combinations to assess their relational proximity in the vector space. We employed the hit ratio at 1 (HR@1) metric, which measures the proportion of cases where the most similar node to a given query node corresponds to a true edge in the KG, treating path-based similarity ranking as a prediction task. A high hit ratio reflects the model's ability to make accurate predictions based solely on embedding similarity. To provide a chance-level reference for evaluating model performance, we also computed random HR@1 baselines for each edge type, under a random ranking assumption. If the top-1 candidate is selected uniformly at random from the $N$ possible nodes, the probability that it corresponds to a true neighbour of a query node $q$ equals the fraction of ground-truth relations it has (i.e., $G_q / N$, where $G_q$ is the number of ground-truth neighbours of $q$, and $N$ is the total number of candidate target nodes). Averaging this value across all query nodes yields the random baseline HR@1 for the edge type.

The results indicated robust performance (Figure 2c), particularly in capturing drug-drug interactions, where the HR@1 is 0.457, meaning that approximately 46% of true drug-drug interaction edges can be found by checking top drug-to-drug embedding similarities. Although the HR@1 scores for other disease- and drug-related edges are lower, the overall results are impressive given the challenge of predicting pairwise relationships solely from embedding similarities. The random HR@1 baselines for all edge types remain very small due to the large size of the candidate node pools, highlighting both the inherent difficulty of the task and the meaningful improvement achieved by the model over chance-level performance. These findings suggest that representations derived from the diverse relational information encoded in CROssBARv2 KG can effectively support the prediction of novel drug and disease associations.

To assess the biological relevance of specific relationships, we examined the drug "Coagulation factor VIIa Recombinant Human" (DB00036) as an illustrative example (Figure 2d). In the KG, this drug is connected to two diseases—Glanzmann thrombasthenia 1 (MONDO:0031332) and haemophilia A (MONDO:0010602)—which were also ranked as the most similar disease nodes in the embedding space, with cosine similarities of 0.909 and 0.805, respectively. Moreover, the top five most similar drugs to DB00036—DB13923, DB11330, DB13151, DB00513, and DB06692—showed high cosine similarities ranging from 0.704 to 0.891, and are all linked to DB00036 via drug-drug interaction edges in the KG. These alignments indicate that the embeddings effectively capture both disease and drug relationships encoded in the KG.

"Antihemophilic factor, human recombinant" (DB00025) ranks as the next most similar drug to DB00036 in the embedding space, with a cosine similarity of 0.659. Unlike the top five most similar drugs to DB00036, which are all connected to it via known drug-drug interactions, DB00025 has no such interaction recorded, making it a compelling candidate for exploring potential novel associations. An examination of DB00025's neighbourhood reveals that it is connected to DB00036 through two different 2-step paths, and haemophilia A and Glanzmann thrombasthenia 1—diseases associated with DB00036—through three-step paths (Figure 2d). This structural proximity validates the high embedding-based similarity between DB00025 and DB00036, and suggests that DB00025 may be related to diseases associated with DB00036 through shared contextual relationships. Notably, DrugBank reports DB00025 as a treatment for haemophilia A, a condition characterised by insufficient production of antihemophilic factor (AHF) (https://go.drugbank.com/drugs/DB00025). The association between haemophilia A and AHF is also well-documented in the literature [31,32], indicating that embedding similarity-based assessments can uncover valid therapeutic indications even in the absence of direct annotations in source databases. Although pathway, side effect, and phenotype nodes—each potentially relevant for uncovering drug- and disease-related associations—were not included in the metapath definitions used for embedding generation in this use case, Figure 2d shows that these entities form indirect connections between the drug and the diseases discussed. Integrating these node types into the embedding process could enrich the resulting representations and enhance the system's capacity to uncover novel therapeutic associations, reinforcing the value of comprehensive data integration for robust biological modelling.

#### S2.3. Assessing cross-database pathway synonymy

Lastly, to assess the biological coherence of pathway relationships in the KG, we analysed curated pairs of synonymous pathways from KEGG and Reactome databases that represent the same metabolic or signalling pathways. We aimed to determine if the metapath2vec embeddings accurately captured pathway content and function—as reflected in their relationships within the KG—by evaluating whether synonymous pathways were positioned closely in the embedding space. Proximal embedding of synonymous pathways would reflect the KG's ability to capture biologically relevant pathway relationships.

We generated the embeddings by defining hierarchical pathway relationships and protein–pathway interactions as metapaths, allowing the representations to capture both pathway organisation and associated molecular features. Cosine similarities were calculated between the embedding vectors, and for each pathway node, a ranked list of other pathway nodes was generated based on their similarity scores. HR@10 metric was then computed based on true KEGG-Reactome synonymous pairs, which yielded a mean score of 0.426. This result suggests that metapath2vec effectively captures synonymous relationships, with approximately 43% of pathway nodes having their curated exact-match counterparts ranked among the top 10 most similar pathways in the embedding space. In this analysis, only strict matches were considered in the ground-truth dataset, excluding highly similar or related but non-identical pathway matches, which are challenging to take into account due to complex hierarchical pathway relationships. If those could be included, the actual rate of biologically meaningful matches would likely have been significantly higher.

To closely examine the synonymous relationships that were captured with high similarity based on representation vectors, we focused on Reactome pathway nodes that had their true synonyms ranked within the top three most similar nodes. Three examples of these are presented in Figure 2e. The results showed that pathway nodes representing identical or highly similar metabolic processes exhibited high similarity scores, often around or above 70%. As the similarity scores decreased, the pathways became less related. For instance, the Reactome pathway for“ acid metabolism” (R-HSA-8978868) showed the highest similarity (0.759) with its KEGG synonym hsa01212. The KEGG pathway for“ acid elongation” (hsa00062), which is associated with the anabolic processes of fatty acid metabolism, also showed a high similarity score (0.735) with R-HSA-8978868. Conversely, the “beta-Alanine metabolism” pathway (hsa00410), which is not directly related to fatty acid metabolism, had a lower similarity (0.549) with R-HSA-8978868. By incorporating protein–pathway associations and pathway hierarchies, metapath2vec captures the contextual features that define functional similarity, allowing it to place synonymous or similar pathways close together in the embedding space. Such integration of diverse biological information supports a more complete understanding of biological systems and helps uncover hidden connections between related processes.

#### S2.4. Elucidating Potential Therapeutic Mechanisms of De Novo Molecules via CROssBARv2’s Hybrid Approach

Here, we aimed to demonstrate the utility of the CROssBARv2 KG in identifying potential therapeutic mechanisms. In this drug discovery-centric scenario, the user seeks to gain a prior understanding of the therapeutic potential and possible mechanism of action of five structurally analogous *de novo* drug-like (query) molecules (**Figure 3f**, left part). These molecules are completely novel entities not present in any biological/chemical databases, with only chemical structures known, which makes this task highly challenging without wet-lab experimentation.

To accomplish this, a hybrid approach combining vector similarity search and subgraph analysis, both part of the CROssBARv2 framework, are utilised. As the first step, the user obtains numerical representations (i.e., embeddings) of his/her small molecules using the SELFormer [33] chemical language model (https://github.com/HUBioDataLab/SELFormer), the same embedding model used in CROssBARv2 for existing drug/compound nodes. After that, the user queries the *de novo* molecule embeddings against our KG using the semantic similarity search option in the CROssBARv2 system to identify relevant biological entities and relationships, resulting in a subgraph that reveals key interactions with existing drugs/compounds, genes/proteins, subcellular locations, organisms, diseases, and side effects (**Figure 3f**).

The vector similarity search demonstrates high cosine similarity (>0.99) between the query molecules and a drug-like compound from the ChEMBL Database (id: CHEMBL4076361), which indicates significant structural resemblance between them. As shown in the resulting graph (**Figure 3f**), CHEMBL4076361 is an inverse agonist of the mouse Cannabinoid receptor 1 protein (UniProt accession: P47746, gene symbol: Cnr1), which has been documented via curated bioassay(s) [34]. Orthology relationships showed that mouse Cnr1 is orthologous to human CNR1 (UniProt accession: P21554), suggesting translational therapeutic relevance. Further exploration revealed that human Cannabinoid receptor 1 and Nitric oxide synthase, inducible (UniProt accession: P35228, gene symbol: NOS2) proteins have been implicated in common metabolic and cardiovascular conditions, including obesity, cardiovascular disorders, and diabetes mellitus, as supported by literature findings [35–42]. The shortest path length between CNR1 and NOS2 proteins in the KG is 3. One such path, proceeding via Guanine nucleotide-binding protein G(i) subunit alpha-2 (UniProt accession: P25311) and Nitric oxide synthase 3 (UniProt accession: P29475), is depicted in the graph (**Figure 3f**). Furthermore, CNR1 and NOS2 proteins have high sequence embedding cosine similarity (0.93), and both enable protein binding and are active in the plasma membrane, indicating potential functional similarities. Regarding their individual functions, CNR1 is involved in regulation of metabolic process and takes part in Retrograde Endocannabinoid Signaling pathway, while NOS2 protein enables NADP binding and participates in Nitric Oxide Stimulates Guanylate Cyclase pathway. These pathways are further modulated by diseases linked to genes encoding CNR1 and NOS2 proteins, highlighting their shared pathological contexts. At the gene level, both CNR1 and NOS2 are transcriptionally activated by JUND, a regulatory factor identified through integrated gene regulatory network resources. Ibipinabant, an anti-obesity drug targeting CNR1 protein, and Nimodipine, a cardiovascular agent targeting both CNR1 and NOS2 proteins, were identified via small molecule-target interactions from bioassay and drug-related databases in our system (e.g., DrugBank, ChEMBL, PHAROS, etc). Notably, Nimodipine and Ibipinabant exhibited significant cosine similarity (>0.98) to CHEMBL4076361 in the vector search, reinforcing their functional relevance. In another observation, CHEMBL4076361 showed significant cosine similarity (>0.99) to Rimonabant, a well-known anti-obesity drug and the inverse agonist of the CNR1 protein that was withdrawn from clinical use due to neuropsychiatric side effects [35–42].

In summary, our hybrid approach has successfully elucidated potential therapeutic mechanisms of *de novo* analogue molecules absent from existing chemical databases. The high semantic similarity (>0.99) between the de novo molecules and CHEMBL4076361 indicates strong structural and functional resemblance. Orthology relationship indicates that findings related to mouse Cnr1 can be extrapolated to the human CNR1, while the high cosine similarity between CHEMBL4076361 and known NOS2-targeting drugs implies that CHEMBL4076361 may also be functionally relevant to NOS2. Together, these connections suggest that the de novo molecules may act as dual modulators of CNR1 and NOS2, and thus could be applicable to conditions such as obesity, cardiovascular disorders, or diabetes mellitus. Despite no direct interaction between CNR1 and NOS2 proteins exists in the KG, their shared disease associations, subcellular localization, molecular function, transcriptional regulation, and high vector similarity collectively point to potential biochemical or functional crosstalk. Overall, this methodology connects uncharacterised biological entities to established biomedical knowledge, yielding valuable insights into the potential biological implications. This methodology is particularly promising in the current era of deep learning-driven molecule generation, where novel biomolecules are being designed at an unprecedented scale. These *de novo* molecules often lack experimental validation or prior annotations, making it challenging to infer their biological roles. Our approach can significantly accelerate the evaluation and prioritisation of these AI-generated molecules, bridging the gap between computational design and biological understanding.

### S3. The CROssBAR-LLM interface

CROssBAR-LLM provides an integrated interface (Figure S1) that enables users to navigate and query the KG using natural language, with the option to leverage precomputed vector embeddings to identify entities based on semantic similarity. For each query, the interface presents a concise natural-language response, followed by a detailed result view in the right panel, which includes:

- Generated Cypher Query: The Cypher query automatically generated by the LLM based on the user's natural language input.
- Node Information: A structured list of nodes retrieved by the generated query, including direct links to their corresponding records in external databases.
- Structured Query Results: The JSON-formatted data returned from the database after execution of the Cypher query.
- In addition, each response is accompanied by three suggested follow-up questions generated by the LLM. Users can continue the analysis either by selecting one of these suggestions or by submitting their own follow-up queries, enabling iterative and progressively refined exploration of the knowledge graph.

**a** Direct graph querying

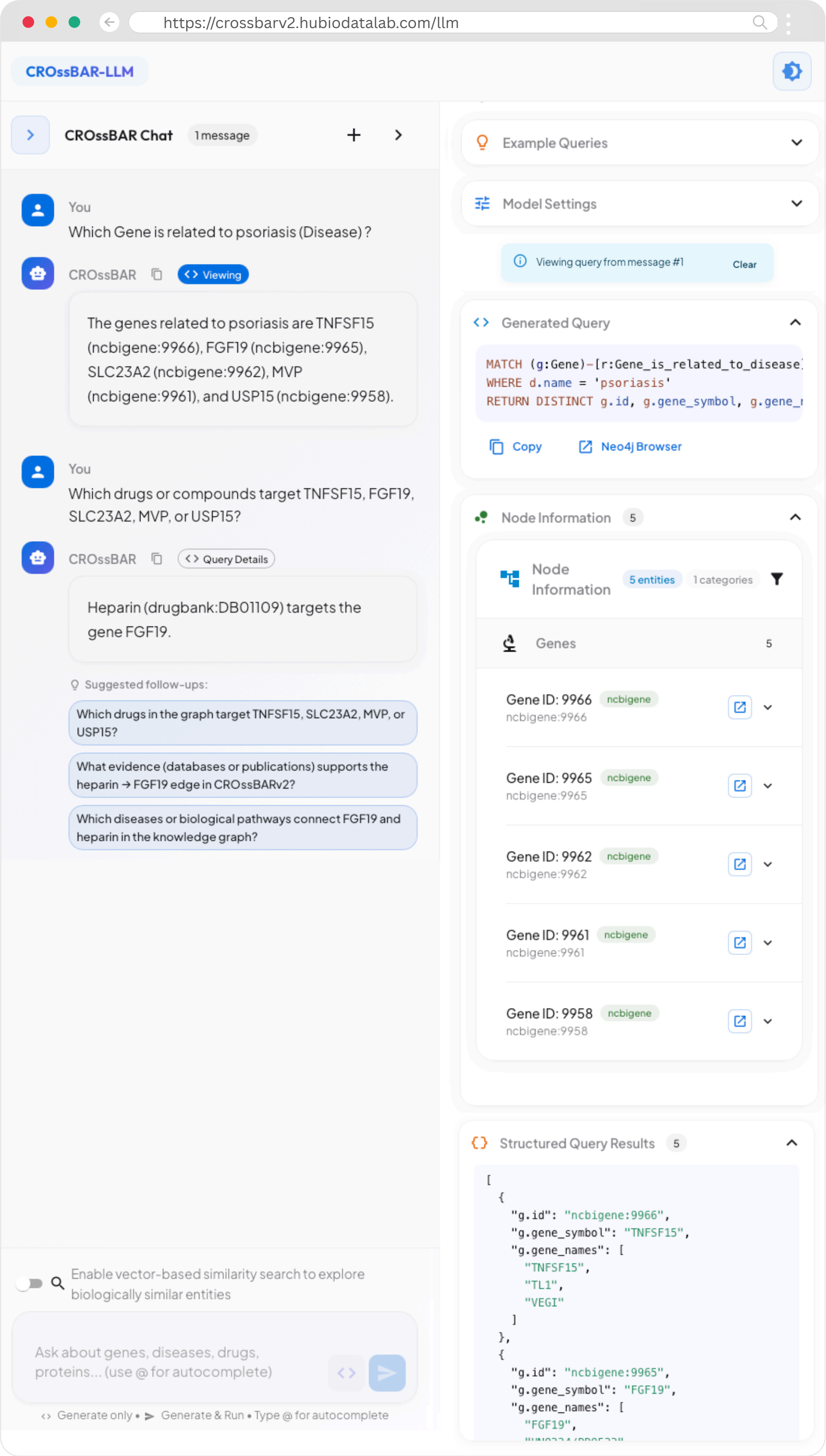

**b** Vector-based similarity search enabled

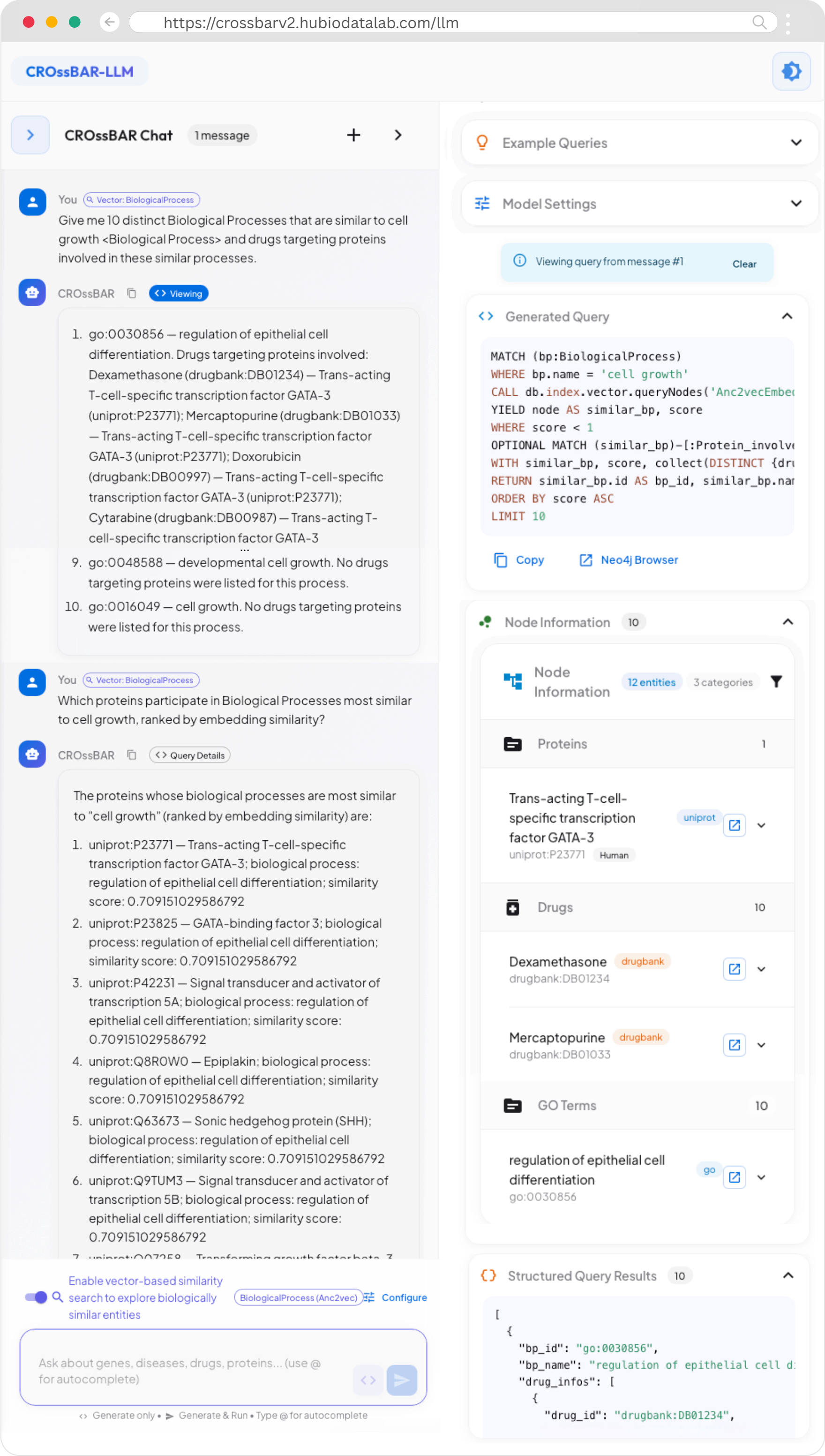


**Figure S1. CROssBAR-LLM interface. (a)** CROssBAR-LLM allows users to interact with the knowledge graph by translating natural-language inputs into Cypher queries and displaying retrieved node information and structured results. **(b)** When vector-based similarity search is enabled, precomputed embeddings are leveraged to retrieve biologically related entities based on semantic similarity in addition to explicit graph connections.

### S4. A qualitative comparison between CROssBAR-LLM and standalone LLMs in complex biomedical question-answering, extended results

S4.1. Query-1

Our first query: "*What drugs are used for 'obesity disorder' disease and have drug interactions with 'Ceritinib', while also targeting proteins involved in the 'ABC transporters' pathway?*", was designed to test a sole LLM's and our system's ability to integrate multiple layers of biological and biomedical knowledge. Answer to this requires combining information on drug-disease relationships, drug-drug interactions, and drug-protein associations tied to a specific biological pathway. The correct set of drugs in the output should interact with ceritinib, a tyrosine kinase inhibitor (TKI) primarily used for treating non-small cell lung cancer (NSCLC) with ALK gene rearrangements, either directly through its impact on liver enzymes, particularly cytochrome P450 (CYP), or indirectly via its association with processes like QT prolongation and hepatic transporters. Another aspect of this query involved ABC transporters, such as ABCB1 (P-glycoprotein), which play a vital role in maintaining cellular drug homeostasis by translocating drugs across cellular membranes to regulate their intracellular concentrations. Ceritinib, as both a substrate and an inhibitor of ABCB1, can disrupt this balance by reducing the efflux of drugs that rely on this transporter. This inhibition can lead to the intracellular accumulation of such drugs, potentially altering their pharmacokinetics, therapeutic efficacy, and toxicity profiles.

When we ran this query through our interface, CROssBAR-LLM retrieved structured data from the knowledge graph (KG), identifying 12 drugs that met the specified criteria based on their documented interactions and pathway involvement. The system generated a Cypher query, executed it on the Neo4j graph database, and retrieved drugs that (i) are indicated for obesity disorder, (ii) interact with Ceritinib, and (iii) target proteins within the ABC transporters pathway. The retrieved drugs are presented in Table S1, providing a structured view of the response.

Among the drugs identified, sertraline, rosuvastatin, and bromocriptine exhibit stronger connections to obesity and its comorbidities. Sertraline is indicated for eating disorders, which frequently co-occur with obesity due to behavioural and psychological factors. Rosuvastatin is prescribed to reduce cholesterol levels, a common metabolic concern in obese individuals, while bromocriptine is used to treat type 2 diabetes mellitus, a condition closely linked to obesity through its effects on glucose metabolism. As visualised in Figure 4a, these drugs maintain key relationships with central entities in the query, including Ceritinib, obesity disorder and the ABC transporters pathway. Apart from that, some output drugs are primarily used in clinical settings for purposes such as sedation and general anaesthesia (e.g., sevoflurane and propofol) or for treating bacterial infections and hormonal disorders (e.g., azithromycin and hydrocortisone). Due to their broad therapeutic applications, these drugs are often associated with multiple diseases in biomedical databases, reflecting their general utility rather than a specific relevance to obesity.

**Table S1.** Formatted version of the response from the CROssBAR-LLM system to the query: "*What drugs are used for obesity disorder and have drug interactions with Ceritinib while also targeting proteins involved in the ABC transporters pathway?*"

| Drug Name | Drug Identifier (DrugBank) |
|---|---|
| Sevoflurane | DB01236 |
| Bromocriptine | DB01200 |
| Pioglitazone | DB01132 |
| Sertraline | DB01104 |
| Rosuvastatin | DB01098 |
| Propofol | DB00818 |
| Fentanyl | DB00813 |
| Hydrocortisone | DB00741 |
| Midazolam | DB00683 |
| Warfarin | DB00682 |
| Fluoxetine | DB00472 |
| Azithromycin | DB00207 |

GPT-4o was prompted to answer the same query using its web search functionality, ensuring access to the most recent information available online. The model returned the following response:

*"Obesity is a complex condition often managed with medications that can interact with other treatments. Ceritinib, a tyrosine kinase inhibitor used in certain lung cancers, is metabolized primarily by the enzyme CYP3A4. Therefore, drugs that inhibit or induce CYP3A4 can alter ceritinib's effectiveness and safety.*

*Several anti-obesity medications are known to interact with ceritinib:*

- *Naltrexone/Bupropion (Contrave, Mysimba): Bupropion is metabolized by CYP2B6 and can inhibit CYP2D6, potentially affecting ceritinib metabolism. Caution is advised when co-administering these drugs. (https://en.wikipedia.org/wiki/Naltrexone/bupropion?utm_source=chatgpt.com)*
- *Bupropion/Zonisamide (Empatic): Although still experimental, this combination includes bupropion, which may interact with ceritinib similarly to the naltrexone/bupropion combination. (https://en.wikipedia.org/wiki/Bupropion/zonisamide?utm_source=chatgpt.com)*

*Regarding the ATP-binding cassette (ABC) transporters, these proteins play a significant role in drug absorption, distribution, and elimination. Alterations in their expression can influence the pharmacokinetics of various medications, including those used in obesity management. However, specific interactions between ceritinib and anti-obesity drugs targeting ABC transporters are not well-documented. (https://www.mdpi.com/2077-0375/14/11/223?utm_source=chatgpt.com)*

*Given the potential for drug interactions, it's crucial to consult with a healthcare professional before combining ceritinib with any anti-obesity medication. They can provide personalised advice based on your medical history and current treatments."*

While it accurately emphasised known anti-obesity drugs with potential interactions with ceritinib due to its metabolism by CYP3A4, its response lacked specificity regarding drug-protein interactions within the ABC transporter pathway. GPT-4o noted the relevance of CYP3A4 metabolism in ceritinib's interactions and the role of ABC transporters in drug pharmacokinetics. However, it acknowledged the limited documentation on specific interactions between ceritinib and anti-obesity drugs targeting ABC transporters, leaving the query only partially addressed. GPT-4o's response is tailored for general readability and is easy to understand for a broad audience. It primarily relies on summarised insights, likely drawn from online resources such as Wikipedia. Consequently, when confronted with a query that cannot be resolved by any existing resource and thus demands deeper reasoning, it fails to produce a correct answer.

In contrast, CROssBAR-LLM delivered a more comprehensive output, identifying a broader set of drugs. By integrating multiple layers of biological data, our tool not only identified the drugs meeting the query's criteria but also highlighted the underlying biological mechanisms driving these interactions, demonstrating its ability to navigate complex biomedical queries with precision.

#### S4.2. Query-2

Our second query: "*What proteins are encoded by genes regulated by 'NFKB1' gene, take part in 'Endocytosis' pathway, and are targeted by drugs used to treat diseases comorbid with 'osteoporosis' disease?*" requires combining insights into gene regulation networks, pathway annotations, drug-target interactions, and disease comorbidity data. To answer it satisfactorily, a tool/model must demonstrate the ability to integrate and interpret this complex information. Specifically, it must first identify proteins encoded by genes regulated by NFKB1 (UniProt accession: P19838), a transcription factor involved in various cellular processes, including inflammation and immune response. Next, it must determine which of these proteins participate in the Endocytosis pathway (KEGG pathway id: hsa04144), a critical cellular mechanism for nutrient uptake, signal transduction, and pathogen entry. Finally, it should intersect this data with drug-target information to identify drugs acting on these proteins that are used to treat diseases comorbid with osteoporosis, such as arthritis, depression, or cardiovascular diseases [43].

The CROssBAR-LLM system extracted structured data from KG, identifying 19 proteins that satisfied the specified criteria based on their regulation, pathway involvement, and drug interactions. A Cypher query was generated and executed on the Neo4j graph database to retrieve proteins that (i) are encoded by genes regulated by NFKB1, (ii) participate in the endocytosis pathway, and (iii) are targeted by drugs used to treat diseases comorbid with osteoporosis. The retrieved proteins are listed in Table S2. Among these output proteins, proto-oncogene tyrosine-protein kinase Src (P12931) was found to be a central player in cellular signalling, mediating pathways linked to cell survival, proliferation, and migration. Additionally, HLA class I histocompatibility antigens (HLA-G, HLA-E, HLA-A) were highlighted for their essential role in antigen presentation and immune response regulation. The Epidermal growth factor receptor (EGFR) (P00533) was identified as a key regulator of cell division and differentiation, primarily through its activation of growth-promoting pathways. Figure 4b visualises how these proteins are linked to key aspects of the query, including regulation by NFKB1, participation in the endocytosis pathway, and targeting by drugs such as imatinib and zonisamide, which are used for diseases comorbid with osteoporosis. Beyond these, the system identified related proteins contributing to immune regulation, inflammatory response, cytoskeletal dynamics, and signal transduction.

**Table S2.** Response from the CROssBAR-LLM system to the query: “What proteins are encoded by gene regulated by 'NFKB1' gene, take part in 'Endocytosis' pathway, and are targeted by drugs used to treat diseases comorbid with 'osteoporosis' disease?”

| Protein Name | Protein Identifier |
|---|---|
| Mothers against decapentaplegic homolog 3 | P84022 |
| Interleukin-2 receptor subunit alpha | P01589 |
| C-X-C chemokine receptor type 1 | P25024 |
| Proto-oncogene tyrosine-protein kinase Src | P12931 |
| Insulin-like growth factor 1 receptor (EC 2.7.10.1) (Insulin-like growth factor I receptor) (IGF-I receptor) (CD antigen CD221) | P08069 |
| Epidermal growth factor receptor | P00533 |
| C-X-C chemokine receptor type 2 | P25025 |
| E3 ubiquitin-protein ligase Mdm2 | Q00987 |
| HLA class I histocompatibility antigen, alpha chain G (HLA G antigen) (MHC class I antigen G) | P17693 |
| HLA class I histocompatibility antigen, alpha chain E (MHC class I antigen E) | P13747 |
| HLA class I histocompatibility antigen, A alpha chain | P04439 |
| Caveolin-1 | Q03135 |
| G protein-coupled receptor kinase 5 | P34947 |
| Cell division control protein 42 homolog | P60953 |
| Beta-arrestin-2 | P32121 |
| C-C chemokine receptor type 5 | P51681 |
| Platelet-derived growth factor receptor alpha | P16234 |
| TGF-beta receptor type-2 | P37173 |
| Protein PML | P29590 |

In comparison, GPT-4o was queried using its web search functionality to access the most up-to-date information, and the resulting response is provided below.

*"Identifying proteins that are regulated by the NFKB1 gene, participate in the endocytosis pathway, and are targeted by drugs used to treat diseases comorbid with osteoporosis involves complex interactions. While specific proteins meeting all these criteria are not readily identifiable from the available data, several relevant connections can be highlighted:*

1. *NFKB1 Gene Regulation: The NFKB1 gene encodes a subunit of the NF-κB protein complex, which is a transcription factor involved in various cellular processes, including immune and inflammatory responses. It regulates the expression of numerous genes, some of which may be involved in endocytosis. (https://www.genecards.org/cgi-bin/carddisp.pl?gene=NFKB1&utm_source=chatgpt.com)*
2. *Endocytosis Pathway: Endocytosis is a cellular process by which cells internalize molecules and particles from their surroundings. While NF-κB signaling can influence various cellular pathways, direct regulation of endocytosis-related proteins by NFKB1 is not well-documented.*
3. *Comorbid Diseases and Drug Targets: Osteoporosis often co-occurs with conditions such as rheumatoid arthritis and certain cancers. Drugs used to treat these conditions may target pathways involving NF-κB signaling. For example, some anti-inflammatory drugs used in rheumatoid arthritis treatment aim to inhibit NF-κB activity to reduce inflammation. (https://www.frontiersin.org/journals/pharmacology/articles/10.3389/fphar.2024.1456796/full?utm_source=chatgpt.com)*

*In summary, while NF-κB signaling is broadly involved in various cellular processes and diseases, pinpointing specific proteins that are regulated by NFKB1, involved in endocytosis, and targeted by drugs for osteoporosis-comorbid diseases requires more detailed research. Consulting specialized databases or scientific literature focusing on protein interactions and signaling pathways may provide more precise information."*

GPT-4o broke the problem into its key components and offered biological context on NFKB1-regulated proteins, the endocytosis pathway, and diseases comorbid with osteoporosis. However, its response lacked the specificity needed to identify proteins that met all the criteria simultaneously, relying instead on general biological knowledge without traceable data from curated sources. Notably, GPT-4o suggested that consulting specialised databases or scientific literature could yield more precise information, a gap addressed by the KG's structured data derived from reliable sources. While GPT-4o excels in readability and contextual framing, CROssBAR-LLM delivers structured and traceable data essential for detailed biomedical investigations.

Together, these use cases illustrate the potential of combining natural language models with knowledge graphs to produce precise, biologically meaningful, and contextually relevant responses to complex biomedical queries. CROssBAR's structured data is directly traceable to curated sources, ensuring reliability and enabling further computational analyses. This systematic approach offered greater depth, transparency, and practical utility for detailed biomedical research and integration into downstream workflows.

### S5. CROssBAR-LLM benchmarks on Cypher query generation and biomedical question answering

In this section, we evaluated the performance of LLMs in two key areas: (i) generating Cypher queries from natural language for structured database interaction and (ii) answering biomedical questions with and without structured knowledge augmentation. In the first one, we assessed how well different LLMs used in CROssBAR-LLM translate natural language prompts into valid Cypher queries, examining the role of model size and computational constraints. This analysis was critical to achieving the optimal performance of CROssBAR-LLM. In the second area, we conducted two tests. We investigated the impact of structured knowledge integration by evaluating LLMs on a true-false-based biomedical question-answering benchmark, comparing standalone LLM performances to the same models augmented with the CROssBARv2 KG (i.e., CROssBAR-LLMs). Then, we test the generalisation ability of CROssBAR-LLMs on an independent genomics dataset, exploring their strengths and limitations compared to web search-augmented standalone LLMs.

#### S5.1. Assessing the Capability of Large Language Models in Generating Cypher Queries from Natural Language

To evaluate how effectively CROssBARv2, backed by different LLMs, can generate valid Cypher queries—a crucial function of CROssBARv2—we constructed a benchmark dataset of 50 natural language questions curated from our graph database, e.g., “Find all drugs that decrease the expression of genes related to 'Duchenne and Becker muscular dystrophy'”, (the complete query dataset is provided in **Supplementary Files**). We tasked these LLMs with generating queries for the curated questions, and then compared their outputs to the ground truth answers. The performance results in **Figure 5a** revealed that open-source models, including DeepSeek R1 (41/50), LLaMA 3.1-405B (39/50) and Mistral Large 2 (41/50), demonstrated performance at the level of the state-of-the-art proprietary models (e.g., Claude 3.5 and GPT-4 series), highlighting their potential in structured query generation task. Another observation is that model performance correlates with parameter size, with larger models generally yielding better results. Additionally, the LLaMA 3 series demonstrate a clear trend of improved performance across versions of the same parameter scale. To evaluate the feasibility of deploying a model under resource-constrained conditions, we tested a quantised version of Llama 3.1-8B deployed on a local computer via the Ollama [44] framework. This lightweight implementation exhibited markedly poor performance, with only 2 correct answers. Although this model is advantageous for deployment on standard computers or servers due to its reduced computational requirements, the results reflect their performance-related limitations. Lastly, to specifically assess performance on more complex graph traversals, we analyzed a subset composed of 11 questions from our benchmark requiring Cypher queries with a path length of 3 or more hops, e.g., “Which side effects are caused by drugs that target proteins involved in the 'chemical synaptic transmission' biological process?”. Evaluations on this complex query subset revealed strong performance from several leading models (e.g., GPT-4 series, and DeepSeek R1 achieving 9/11 correct answers), where smaller models mostly struggled (e.g., Llama 3 70B, and GPT-4o-mini scored 2/11 correct answers). These results are consistent with the overall trends observed in the full 50-question benchmark, where larger and more advanced models generally demonstrated superior performance.

Overall, our empirical analysis reveals that state-of-the-art LLMs demonstrate considerable proficiency in generating Cypher queries, showcasing their advanced natural language understanding and translation capabilities. These results indicate that CROssBAR-LLM, when backed by one of these top models, can generate a valid query 84% of the time—an outcome that can be deemed satisfactory.

#### S5.2. Integration of CROssBARv2 KG Significantly Enhances LLM Accuracy in Biomedical Question Answering

We developed a benchmark dataset comprising 100 simple true-false questions, e.g., "AGER Gene is associated with acute monocytic leukemia. Answer only True or False!", (the complete question set is given in **Supplementary Files**) to assess the performance of LLMs in answering straightforward biomedical queries. These questions are used to evaluate LLMs augmented with the CROssBARv2 KG and to compare the performance with the standalone LLMs. We also assessed the performance of Sonar Reasoning [45], a web search augmented LLM designed for improved reasoning, to determine whether external web searches could yield performance comparable to CROssBAR-LLMs. We compared the LLM outputs to the ground truth answers for performance evaluation. Whenever CROssBAR-LLMs retrieved no results from the KG, we instructed the underlying LLM to rely on its internal knowledge to generate an answer, thereby allowing for an end-to-end comparison between CROssBAR-LLMs and standalone LLMs and assessing the added value of integrating the CROssBARv2 KG into an LLM-based system.

The correct number of answers for each standalone LLM and CROssBAR-LLM is shown in Figure 5b. Interestingly, our binomial test-based analysis shows that, when answering true-false biomedical questions, most standalone LLMs do not perform significantly better than random guessing (Figure 5b). While Sonar Reasoning outperformed standalone LLMs, achieving an accuracy of 71%, its performance remained considerably lower than all CROssBAR-LLMs. Overall, augmenting LLMs with biomedical knowledge from CROssBARv2 KG led to a substantial performance boost, with models such as GPT-4o (97%) and Gemini 1.5 Pro (98%) achieving near-perfect results. These findings underscore the critical importance of domain-specific knowledge integration to enhance the reliability and accuracy of LLMs for biomedical question answering. Furthermore, these results suggest that in the biomedical domain, where crucial information is dispersed across numerous data sources, structurally organised knowledge, as in CROssBARv2, provides a more consolidated and contextually grounded approach for augmenting LLMs than general web search strategies.

#### S5.3. Generalisation Capabilities of CROssBAR-LLMs on Externally Curated Genomic Benchmark

To evaluate the capabilities of CROssBAR-LLMs and web search-augmented LLMs beyond our in-house benchmarks, we employed the independently curated GeneTuring benchmark [46]. This choice allows for a fair comparison, providing an unbiased assessment of each model's ability to generalise beyond our KG's scope. GeneTuring is a comprehensive benchmark for genomics-focused LLM performance and comprises multiple modules. Within the scope of CROssBARv2, we selected the "transcription factor regulation" dataset composed of 50 binary selection questions as activation/repression, e.g., "Does transcription factor MSC activate or repress gene CDC6?", (the complete question set is given in **Supplementary Files**)

Our evaluation demonstrated that CROssBAR-LLMs generally outperformed standalone LLMs utilising web search (Figure 5c). Claude 3.5 Sonnet and DeepSeek R1 backed CROssBAR-LLMs achieved the highest accuracy (43 and 45 correct answers, respectively), whereas web search-augmented standalone GPT-4o and Gemini-1.5-Pro performed slightly worse (39 and 38 correct answers, respectively). Interestingly, deviating from the general trend, Gemini-1.5-Pro with web search outperformed its CROssBAR counterpart (with 38 to 34 correct answers, respectively). Although Gemini-1.5-Pro and LLaMA 3.1-405B demonstrated an understanding of converting semantically equivalent questions into Cypher queries, their execution was inconsistent, occasionally generating hallucinated or erroneous queries that led to retrieval failures or incorrect responses. The overall performance difference suggests that, while web search offers a broad mechanism for information retrieval, direct access to structured, domain-specific knowledge—such as that provided by CROssBARv2—yields more precise and reliable responses, provided that consistent database queries can be generated.

### S6. Deep learning-based high-performance prediction of biological relationships using CROssBARv2

Protein function prediction is a key task in biomedical research, crucial for understanding cellular processes, disease mechanisms, and identifying potential therapeutic targets. Heterogeneous knowledge graphs provide a robust framework for this task by integrating data across multiple biological domains, enabling deep learning models to learn rich, context-aware representations. To evaluate the effectiveness of the CROssBARv2 KG in this context, we developed a model called ProtHGT [47], which formulates protein function prediction as a link prediction task between protein and Gene Ontology (GO) term nodes (Figure 6a). Using initial node feature vectors, the Heterogeneous Graph Transformer (HGT)-based model updates node representations through multiple layers of type-aware message passing, allowing each node to integrate information from its graph neighbourhood. The final embeddings of each protein–GO term pair are concatenated and passed through a multi-layer perceptron (MLP), followed by a sigmoid activation function to predict the probability of the functional association.

We implemented the model using HGTConv layers from PyTorch Geometric [48] and trained separate models for each GO sub-ontology (i.e., biological process, cellular component, molecular function). Initial node features followed the representations incorporated in the CROssBARv2 KG (see "Building the Knowledge Graph" section). For proteins, we experimented with ProtT5 and ESM2 embeddings. Model performance was evaluated using the time-based DeepHGAT [49] benchmark dataset, which uses training annotations prior to September 1, 2021 and test annotations curated between September 1, 2021 and April 1, 2023. To ensure fair comparison, we constructed time-aligned versions of the CROssBARv2 KG by removing all protein–GO term edges and re-adding only those corresponding to the respective benchmark training and test periods. These time-specific graphs were then used for independent model training, optimisation, and evaluation. Model performance was compared directly against results reported for other methods in the original benchmark publication.

The DeepHGAT benchmark covers a broad range of prediction approaches, including heterogeneous graph-based models [49], protein-protein interaction (PPI)-based models [50–54], and sequence-based approaches [55–58], which rely on pre-trained protein language embeddings. ProtHGT achieved the best performance across all GO sub-ontologies (**Figure 6b**), outperforming even the strongest competitors such as DeepHGAT [49]—which incorporates PPI networks, GO structure, and sequence features—and PSPGO [54], a GNN-based model operating on PPI and sequence similarity networks. This improvement is primarily attributed to ProtHGT's use of the CROssBARv2 KG, which integrates a wide spectrum of biological knowledge, including pathways, domains, and ontological associations—critical for capturing the full functional context of proteins. By embedding this diverse information, ProtHGT learns more expressive and biologically meaningful representations. In contrast, models constrained to sequence features or limited edge types are unable to fully capture the complexity of protein function, resulting in reduced predictive performance.

Beyond predictive performance, interpretability is a critical consideration in evaluating protein function prediction models. ProtHGT leverages the structure of a heterogeneous knowledge graph, which not only improves accuracy but also enhances interpretability. By tracing predictions through multi-relational connections within the KG, researchers can better understand the biological rationale behind each output. These findings emphasise the value of heterogeneous data integration for accurate and interpretable protein function prediction.

### S7. Supplementary Discussion

#### S7.1. Broader Applications of Semantic Similarity in CROssBARv2

Existing approaches to semantic similarity search remain narrowly focused on protein-centric tasks, such as homology detection and functional annotation, leaving broader biological entities and relational contexts underexplored [59–62]. CROssBARv2 addresses this gap by systematically integrating embeddings from diverse deep learning models across biological entities, establishing a unified framework for exploring semantic relationships across multiple biological domains. By extending semantic search beyond protein-centric tasks, this integration significantly enhances the system's utility and discovery potential for complex biomedical inquiries. However, it is important to acknowledge that the efficacy of semantic similarity search hinges on the quality of the underlying embedding model's ability to capture semantically meaningful representations.

#### S7.2. Challenges in Data Integration and Conceptual Granularity in CROssBARv2

The first challenge stems from the incomplete nature of many cross-referencing systems. Despite the rigorous curation efforts, cross-references that are provided by these sources can be incomplete. Such gaps can lead to the unwitting creation of redundant or duplicate entities within the KG. This limitation is particularly evident in our pathway nodes, where we incorporated data from both Reactome and KEGG databases owing to their widespread adoption in life sciences research. Although resources like ComPath [63] offer curated pathway mappings between these databases, their coverage remains incomplete. Future efforts to improve these mappings could benefit from leveraging LLMs, which have the potential to accelerate and enhance the curation of comprehensive pathway mappings by automating the identification of semantically equivalent biological processes across databases.

The second challenge centres on managing conceptual granularity. For example, the existence of numerous disease classification systems (e.g., ICD, MeSH, UMLS, SNOMED-CT, DO), each with its own focus and scope, leads to granularity-related differences, where a disease may appear as a single broad concept in one source but as multiple, more specific sub-entities in another. A notable example of this is Alzheimer disease: it is represented by 19 distinct concepts in DO, 27 in SNOMED-CT, and 17 in MeSH. Similar discrepancies have been observed for other disease concepts in previous studies [2,64]. Such discrepancies reflect broader inconsistencies and significantly complicate data integration across disease sources. Addressing this challenge, Mondo provides a unified and interoperable ontology that systematically reconciles these disparate classifications. These considerations motivated our selection of Mondo over the plethora of available disease classification systems in CROssBARv2 KG. While Mondo's disease representations provide substantial value for data integration and knowledge standardisation, its application in clinical settings may not always translate into practical diagnostic utility. For example, Alzheimer's disease is represented by 21 subterms in Mondo. However, it is highly unlikely that a clinician would utilise the full spectrum of Mondo by diagnosing a patient using specific Mondo sub-terms. These fine-grained terms often represent research-based classifications or specific genetic variants not commonly utilised in clinical settings. To mitigate this disconnect between ontological granularity and clinical practice, a potential solution is to adopt PrimeKG's [2] approach of creating supernodes by clustering disease nodes based on the semantic similarity of their names, effectively grouping related disease terms into broader, more general disease concepts.

S7.3. Methodological Alternatives and Design Rationale for the Text-to-Cypher Approach

A prominent alternative to our Text-to-Cypher approach is based on RAG, where structural graph data is first textualised and then retrieved and processed by LLMs. For instance, Soman et al. [65] adopted this method by converting the one-hop neighbourhood of disease entities into sentences and using semantic retrieval to answer biomedical questions. While this method could be beneficial, it presents certain challenges in the context of complex, interconnected biomedical data. Firstly, traditional RAG strategies struggle to capture the intricate structure of KGs because they depend on semantic similarity alone, rather than leveraging the actual connections within the graph [66] . Secondly, the process of textualising a graph for an LLM is inherently problematic and lossy, as the model does not natively comprehend graph structures and can learn spurious correlations, such as assuming a node's high mention frequency indicates its structural importance [67] . In contrast, by dynamically generating Cypher queries, our approach preserves the native graph structure, enabling precise and structurally-aware information retrieval without information degradation inherent in classical RAG pipelines. Nevertheless, it is important to acknowledge that the efficacy of our method is closely tied to the underlying capabilities of LLMs, particularly their ability to generate accurate Cypher queries and to interpret the given KG schema.

### S8. CROssBARv2 data sources

**Tables S3 and S4** present a comprehensive overview of the data sources used for each biological entity and relationship incorporated into CROssBARv2. In this section, we provide detailed information about these sources, including their scope, content, and integration strategy.

UniProtKB is a widely used repository for high-quality protein sequence and functional information [68]. We utilised the curated UniProtKB/Swiss-Prot database for introducing protein nodes to the system, while gene nodes are created using Entrez identifiers, and organism nodes are established based on NCBI taxonomy identifiers associated with these entries. For protein-protein interactions, STRING [69], IntAct [70], and BioGRID [71] databases are integrated into CROssBARv2, each providing essential details such as confidence scores, interaction types (physical/genetic), and experimental methods validating these interactions.

Gene Ontology (GO) offers a structured vocabulary for annotating genes and gene products, organised into three sub-ontologies: molecular function, biological process, and cellular component [72]. Each sub-ontology is represented as a distinct node type in CROssBARv2, with hierarchical relationships among these nodes to reflect the structural organisation within each ontology. GO nodes also connect to proteins and other biological entities, enabling a comprehensive view of functional relationships across diverse biological components. Enzyme Commission (EC) numbers, sourced from the Expasy ENZYME database [73], categorise enzymes based on the chemical reactions they catalyse. By integrating EC nodes, hierarchical EC relationships, and protein-EC relationships, the system effectively represents enzymatic functions, enriching insights into biochemical pathways.

OMA (Orthologous MAtrix) provides evolutionary relationships across 2,326 genomes, capturing pairwise and group orthologs, functional annotations, and synteny data [74]. Due to the extensive orthology data, CROssBARv2 incorporates orthology relationships only for selected organisms frequently studied, balancing depth and scalability. InterPro, a comprehensive resource on protein families and functional regions, incorporates protein signatures from 13 databases in the InterPro Consortium [75]. InterPro data contributes domain nodes and relationships, essential for capturing functional domains that define protein families and their roles.

For drug data, DrugBank serves as the primary source, providing detailed information on FDA-approved and experimental drugs, including targets, effects, and interactions [76]. ChEMBL, a widely recognised resource in drug discovery, contains extensive information on compounds, bioactivity, and target proteins [77]. ChEMBL adds compound nodes and relationships within CROssBARv2, valuable for drug development and bioactivity research. STITCH complements drug-target data with compound-protein relationships derived from biomedical databases and literature, supported by confidence scores from various sources [78]. DGIdb further enriches drug-gene interactions using sources such as publications and online databases [79]. Drug side effect data are gathered from SIDER, OffSIDES, and ADReCS to capture a broad spectrum of drug-adverse reaction (ADR) information. SIDER provides FDA-registered ADRs [80], OffSIDES catalogs non-FDA-listed side effects [81], and ADReCS includes hierarchical ADR classifications [82]. Different identifier systems (ATC,

UMLS, MedDRA) from SIDER support cross-database mapping for comprehensive ADR representation.

The Mondo Disease Ontology (Mondo) offers a community-curated disease vocabulary, ensuring comprehensive and regularly updated disease definitions sourced from external disease databases [83]. Mondo defines disease nodes, while DisGeNET supplies gene-disease and disease-disease relationships along with variant data incorporated as attributes of gene-disease links, contributing depth to these associations [84]. DISEASES extracts disease-gene relationships via text mining and expert-curated databases, assigning confidence scores to these relationships, enriching the disease-gene data in the system [85]. ClinVar archives human genetic variants associated with diseases, supplemented by supporting evidence from various contributors [86], enriching variant attributes in gene-disease relationships. Additionally, the humsavar dataset from UniProtKB/Swiss-Prot adds manually curated polymorphisms and disease mutations, further contributing to gene-disease associations [68].

The Human Phenotype Ontology (HPO) compiles phenotype terms and their associations with diseases from sources like OMIM, Orphanet, and DECIPHER, providing phenotype-disease, phenotype-gene, and phenotype hierarchy relationships, all of which are included in the system [87]. OpenTargets integrates genetic, somatic mutation, expression analysis, drug study, and animal model data to identify and prioritize potential therapeutic drug targets [88]. This resource contributed to disease-gene relationship data within CROssBARv2. PathoPhenoDB catalogs relationships between pathogens and the diseases and phenotypes they cause [89]. While primarily focused on pathogen-phenotype links, PathoPhenoDB's pathogen-disease relationships derived through text mining are also included in the knowledge graph.

The Kyoto Encyclopedia of Genes and Genomes (KEGG) integrates multiple databases organized around pathways, genomics, and biomedical information [90]. This comprehensive resource provided diverse relationships, including gene/protein-pathway, pathway-drug, and pathway nodes, with KEGG's ID conversion system ensuring consistent entity identification. Reactome, a curated pathway database covering processes like metabolism, signaling, and disease mechanisms. It enriches pathway nodes and contributes hierarchical and orthology relationships between pathways, as well as protein-pathway, drug-pathway, and disease-pathway interactions, complete with supporting evidence.

DoRothEA [91], CollecTRI [92], and TRRUST [93] provided transcription factor (TF)-target gene interactions for CROssBARv2, aggregating data from ChIP-seq, perturbation experiments, gene expression predictions, and curated literature. The Target Central Resource Database (TCRD) is developed as part of the Illuminating the Druggable Genome (IDG) initiative to expand knowledge on poorly understood proteins [94]. Pharos, TCRD's web platform, integrates data from 25 additional sources, enriching the system's orthology and drug-target protein data.

The Comparative Toxicogenomics Database (CTD) curates data on chemical exposures and their biological effects, enhancing drug-gene, disease-pathway, and drug-disease relationships in CROssBARv2 [95]. For gene expression, we used Expression Atlas and the

GTEx Portal. Expression Atlas provides curated gene expression data from over 3,000 experiments across 40 organisms [96], while GTEx provides single-cell gene expression data for human tissues and cells, supporting tissue-specific studies [97].

**Table S3.** Biological entity (node) types within the CROssBARv2 KG.

| Node type | Identifier | Source database |
|---|---|---|
| Protein | UniProt | UniProt |
| Gene | Entrez | UniProt |
| Organism (Taxon) | NCBI Tax ID | UniProt |
| Drug | Drugbank | Drugbank |
| Compound | ChEMBL | ChEMBL |
| Side effect | Meddra | SIDER, OffSIDES, ADReCS |
| Disease | MONDO | MONDO |
| Phenotype | HPO | HPO |
| Protein Domain | InterPro | InterPro |
| Pathway | KEGG, Reactome | KEGG, Reactome |
| EC number | EC number | Expasy ENZYME |
| Molecular function | GO | GOA |
| Cellular component | GO | GOA |
| Biological process | GO | GOA |

**Table S4.** Biological relationship (edge) types within the CROssBARv2 KG.

| Source node type | Relationship | Target node type | Source node identifier | Target node identifier | Source database | Filtering thresholds |
|---|---|---|---|---|---|---|
| Protein | Interacts with | Protein | UniProt | UniProt | STRING, BioGRID, IntAct | Only high-confidence interactions (score >0.7) from STRING |
| Protein | Has | Protein Domain | UniProt | InterPro | InterPro | - |
| Protein | Enables / Contributes to | Molecular Function | UniProt | GO | GOA | Electronic annotations (i.e., IEA) filtered out |
| Protein | Located in / Is active in / Part of | Cellular component | UniProt | GO | GOA | Electronic annotations (i.e., IEA) filtered out |
| Protein | Involved in | Biological process | UniProt | GO | GOA | Electronic annotations (i.e., IEA) filtered out |
| Protein | Take part in | Pathway | UniProt | KEGG, Reactome | Reactome | Electronic annotations (i.e., IEA) from Reactome filtered out |
| Protein | Catalyzes | EC Number | UniProt | EC Number | Expasy ENZYME | - |
| Protein | Belongs to | Organism Taxon | UniProt | NCBI Tax ID | UniProt | - |
| Protein | Is associated with | Phenotype | UniProt | HPO | HPO | - |
| Gene | Encodes | Protein | Entrez | UniProt | UniProt | - |
| Gene | Is ortholog to | Gene | Entrez | Entrez | OMA, Pharos | Orthologs between human and model organisms |

| | | | | | | |
|---|---|---|---|---|---|---|
| Gene | Regulates | Gene | Entrez | Entrez | DoRothEA, TRRUST, CollecTRI | Conflicting regulatory modes (i.e., activation, repression, and unknown or activation and repression) were excluded |
| Gene | Is related to | Disease | Entrez | MONDO | ClinVar, DisGeNET, KEGG, DISEASES, humsavar, OpenTargets | - |
| Drug | Interacts with | Drug | Drugbank | Drugbank | DDInter, KEGG | - |
| Drug | Targets | Protein | Drugbank | UniProt | Drugbank, Pharos, ChEMBL, DGIdb, KEGG, STITCH | - |
| Drug | Upregulates / Downregulates | Gene | Drugbank | Entrez | CTD | Contradictory expression annotations (i.e., both increase and decrease expression) were excluded |
| Drug | Has | Side effect | Drugbank | Meddra | SIDER, OffSIDES, ADReCS | - |
| Drug | Has target in | Pathway | Drugbank | KEGG | KEGG, Reactome | - |
| Compound | Targets | Protein | ChEMBL | UniProt | ChEMBL, STITCH | - |
| Disease | Is comorbid with | Disease | MONDO | MONDO | Malacards | - |
| Disease | Is associated with | Disease | MONDO | MONDO | DisGeNET | - |
| Disease | Is a | Disease | MONDO | MONDO | MONDO | - |
| Disease | Is associated with | Phenotype | MONDO | HPO | HPO | Electronic annotations (i.e., IEA) filtered out |
| Disease | Modulates | Pathway | MONDO | KEGG, Reactome | KEGG, CTD | - |
| Disease | Is treated by | Drug | MONDO | Drugbank | ChEMBL, KEGG, CTD | - |
| Disease | Caused by | Organism Taxon (pathogen) | MONDO | NCBI Tax ID | PathoPheno DB | - |
| Protein Domain | Enables | Molecular function | InterPro | GO | InterPro2GO | - |
| Protein Domain | Localizes to | Cellular component | InterPro | GO | InterPro2GO | - |
| Protein Domain | Is involved in | Biological process | InterPro | GO | InterPro2GO | - |
| Molecular function/ Cellular component/ Biological process | Is a / Part of / Negatively regulates / Positively regulates | Molecular function/ Cellular component/ Biological process | GO | GO | GOA | - |
| Phenotype | Is a | Phenotype | HPO | HPO | HPO | - |
| Pathway | Is part of / Is equivalent to / Participates | Pathway | KEGG, Reactome | KEGG, Reactome | Reactome, Compath | - |
| Pathway | Is ortholog to | Pathway | KEGG, Reactome | KEGG, Reactome | KEGG, Reactome | Only between human and other organisms |
| Side effect | Is a | Side effect | Meddra | Meddra | ADReCS | - |
| EC Number | Is a | EC Number | EC number | EC number | Expasy ENZYME | - |

**Table S5.** Embedding models used for node types represented in the CROssBARv2 KG.

| Node type | Embedding model | Input feature | Embedding size |
|---|---|---|---|
| Protein | ProtT5 [98], ESM2 [99] | Protein sequence | 1024, 1280 |
| Gene | Nucleotide Transformer [100] | Nucleotide sequence | 2560 |
| Drug/Compound | SELFormer [33] | SELFIES | 768 |
| Disease | doc2vec [101] | Disease description | 100 |
| Phenotype | CADA [102] | Gene-phenotype relationship network | 160 |
| Protein Domain | dom2vec [103] | Protein sequence | 50 |
| Pathway | BioKEEN [104] | Gene-pathway association network | 200 |
| EC number | RXNFP [105] | Reaction SMILES | 256 |
| Molecular function/Cellular component/Biological process | anc2vec [106] | Gene Ontology Directed Acyclic Graph | 200 |

### Supplementary References


1. Doğan, T. *et al.* CROssBAR: comprehensive resource of biomedical relations with knowledge graph representations. *Nucleic Acids Res.* **49**, e96–e96 (2021).
2. Chandak, P., Huang, K. & Zitnik, M. Building a knowledge graph to enable precision medicine. *Scientific Data* **10**, 1–16 (2023).
3. Lobentanzer, S. *et al.* Democratizing knowledge representation with BioCypher. *Nat. Biotechnol.* **41**, 1056–1059 (2023).
4. Himmelstein, D. S. *et al.* Systematic integration of biomedical knowledge prioritizes drugs for repurposing. *eLife* **6**, (2017).
5. Morris, J. H. *et al.* The scalable precision medicine open knowledge engine (SPOKE): a massive knowledge graph of biomedical information. *Bioinformatics* **39**, btad080 (2023).
6. Fernández-Torras, A., Duran-Frigola, M., Bertoni, M., Locatelli, M. & Aloy, P. Integrating and formatting biomedical data as pre-calculated knowledge graph embeddings in the Bioteque. *Nat Commun* **13**, 5304 (2022).
7. Santos, A. *et al.* A knowledge graph to interpret clinical proteomics data. *Nat. Biotechnol.* **40**, 692–692 (2022).
8. Ioannidis, V. N., Song, X., Manchanda, S., Li, M. & Pan, X. [ALINTI] Drkg-drug repurposing knowledge graph for covid-19. (2020).
9. Zheng, S. *et al.* PharmKG: a dedicated knowledge graph benchmark for bomedical data mining. *Brief. Bioinform.* **22**, (2021).
10. Quan, X., Cai, W., Xi, C., Wang, C. & Yan, L. AIMedGraph: a comprehensive multi-relational knowledge graph for precision medicine. *Database (Oxford)* **2023**, (2023).
11. Vlietstra, W. J., Vos, R., van Mulligen, E. M., Jenster, G. W. & Kors, J. A. Identifying genes targeted by disease-associated non-coding SNPs with a protein knowledge graph. *PLoS One* **17**, e0271395 (2022).
12. Gualdi, F., Oliva, B. & Piñero, J. Predicting gene disease associations with knowledge graph embeddings for diseases with curtailed information. *NAR Genom. Bioinform.* **6**, lqae049 (2024).
13. Gema, A. P. *et al.* Knowledge graph embeddings in the biomedical domain: Are they useful? A

look at link prediction, rule learning, and downstream polypharmacy tasks. *arXiv [cs.LG]* (2023) doi:10.48550/ARXIV.2305.19979.

14. Vilela, J. *et al.* Biomedical knowledge graph embeddings for personalized medicine: Predicting disease-gene associations. *Expert Syst.* **40**, (2023).
15. Nazi, Z. A. & Peng, W. Large language models in healthcare and medical domain: A review. *Informatics (MDPI)* **11**, 57 (2024).
16. Cascella, M., Montomoli, J., Bellini, V. & Bignami, E. Evaluating the Feasibility of ChatGPT in Healthcare: An Analysis of Multiple Clinical and Research Scenarios. *J Med Syst* **47**, 33 (2023).
17. Tian, S. *et al.* Opportunities and challenges for ChatGPT and large language models in biomedicine and health. *Brief Bioinform* **25**, (2023).
18. Jahan, I., Laskar, M. T. R., Peng, C. & Huang, J. A comprehensive evaluation of Large Language Models on benchmark biomedical text processing tasks. *arXiv [cs.CL]* (2023) doi:10.48550/ARXIV.2310.04270.
19. Wang, J. *et al.* On the robustness of ChatGPT: An adversarial and out-of-distribution perspective. *arXiv [cs.AI]* (2023) doi:10.48550/ARXIV.2302.12095.
20. Sun, K., Xu, Y. E., Zha, H., Liu, Y. & Dong, X. L. Head-to-Tail: How Knowledgeable are Large Language Models (LLMs)? A.K.A. Will LLMs Replace Knowledge Graphs? (2023).
21. Ji, Z. *et al.* Survey of hallucination in natural Language Generation. *ACM Comput. Surv.* (2022) doi:10.1145/3571730.
22. Alkaissi, H. & McFarlane, S. I. Artificial Hallucinations in ChatGPT: Implications in Scientific Writing. *Cureus* **15**, e35179 (2023).
23. Bang, Y. *et al.* A multitask, multilingual, multimodal evaluation of ChatGPT on reasoning, hallucination, and interactivity. *arXiv [cs.CL]* (2023) doi:10.48550/ARXIV.2302.04023.
24. Sen, P., Mavadia, S. & Saffari, A. Knowledge graph-augmented language models for complex question answering. in *Proceedings of the 1st Workshop on Natural Language Reasoning and Structured Explanations (NLRSE)* (Association for Computational Linguistics, Stroudsburg, PA, USA, 2023). doi:10.18653/v1/2023.nlrse-1.1.
25. Pan, S. *et al.* Unifying large language models and knowledge graphs: A roadmap. *IEEE Trans. Knowl. Data Eng.* **36**, 3580–3599 (2024).
26. Yang, L., Chen, H., Li, Z., Ding, X. & Wu, X. Give us the facts: Enhancing large language models with knowledge graphs for fact-aware language modeling. *IEEE Trans. Knowl. Data Eng.* **36**, 3091–3110 (2024).
27. Wu, Y. *et al.* Retrieve-rewrite-answer: A KG-to-Text enhanced LLMs framework for knowledge graph question answering. *arXiv [cs.CL]* (2023) doi:10.48550/ARXIV.2309.11206.
28. Sun, J. *et al.* Think-on-Graph: Deep and responsible reasoning of large language model on knowledge Graph. *arXiv [cs.CL]* (2023) doi:10.48550/ARXIV.2307.07697.
29. Van der Maaten, L. & Hinton, G. Visualizing data using t-SNE. *J. Mach. Learn. Res.* **9**, (2008).
30. Atas Guvenilir, H. & Doğan, T. How to approach machine learning-based prediction of drug/compound–target interactions. *Journal of Cheminformatics* **15**, 16 (2023).
31. Schwartz, R. S. *et al.* Human recombinant DNA-derived antihemophilic factor (factor VIII) in the treatment of hemophilia A. recombinant Factor VIII Study Group. *N. Engl. J. Med.* **323**, 1800–1805 (1990).
32. Lusher, J. *et al.* Human recombinant DNA-derived antihemophilic factor in the treatment of

previously untreated patients with hemophilia A: final report on a hallmark clinical investigation. *Journal of thrombosis and haemostasis : JTH* **2**, (2004).

33. Yüksel, A., Ulusoy, E., Ünlü, A. & Doğan, T. SELFormer: Molecular Representation Learning via SELFIES Language Models. *Machine Learning: Science and Technology* **4**, 025035 (2023).
34. Iyer, M. R. *et al.* Design, synthesis, and biological evaluation of novel, non-brain-penetrant, hybrid cannabinoid CB1R inverse agonist/inducible nitric oxide synthase (iNOS) inhibitors for the treatment of liver fibrosis. *J. Med. Chem.* **60**, 1126–1141 (2017).
35. Engeli, S. *et al.* Activation of the peripheral endocannabinoid system in human obesity. *Diabetes* **54**, 2838–2843 (2005).
36. Ghasemi-Gojani, E., Kovalchuk, I. & Kovalchuk, O. Cannabinoids and terpenes for diabetes mellitus and its complications: from mechanisms to new therapies. *Trends Endocrinol. Metab.* **33**, 828–849 (2022).
37. Barutta, F., Mastrocola, R., Bellini, S., Bruno, G. & Gruden, G. Cannabinoid receptors in diabetic kidney disease. *Curr. Diab. Rep.* **18**, 9 (2018).
38. Mukhopadhyay, P. *et al.* CB1 cannabinoid receptors promote oxidative stress and cell death in murine models of doxorubicin-induced cardiomyopathy and in human cardiomyocytes. *Cardiovasc. Res.* **85**, 773–784 (2010).
39. Rajesh, M. *et al.* Cannabinoid 1 receptor promotes cardiac dysfunction, oxidative stress, inflammation, and fibrosis in diabetic cardiomyopathy. *Diabetes* **61**, 716–727 (2012).
40. Noronha, B. T., Li, J.-M., Wheatcroft, S. B., Shah, A. M. & Kearney, M. T. Inducible nitric oxide synthase has divergent effects on vascular and metabolic function in obesity. *Diabetes* **54**, 1082–1089 (2005).
41. Król, M. & Kepinska, M. Human nitric oxide synthase-its functions, polymorphisms, and inhibitors in the context of inflammation, diabetes and cardiovascular diseases. *Int. J. Mol. Sci.* **22**, 56 (2020).
42. Mungrue, I. N. *et al.* Cardiomyocyte overexpression of iNOS in mice results in peroxynitrite generation, heart block, and sudden death. *J. Clin. Invest.* **109**, 735–743 (2002).
43. Puth, M.-T., Klaschik, M., Schmid, M., Weckbecker, K. & Münster, E. Prevalence and comorbidity of osteoporosis- a cross-sectional analysis on 10,660 adults aged 50 years and older in Germany. *BMC Musculoskelet Disord* **19**, 144 (2018).
44. Ollama. https://ollama.com/.
45. Sonar by Perplexity. https://sonar.perplexity.ai/ (2025).
46. Hou, W., Shang, X. & Ji, Z. Benchmarking large language models for genomic knowledge with GeneTuring. *bioRxivorg* (2025) doi:10.1101/2023.03.11.532238.
47. Ulusoy, E. & Doğan, T. ProtHGT: Heterogeneous graph transformers for automated protein function prediction using biological knowledge graphs and language models. *bioRxiv* (2025) doi:10.1101/2025.04.19.649272.
48. Fey, M. & Lenssen, J. E. Fast Graph Representation Learning with PyTorch Geometric. (2019).
49. Zhao, Y. *et al.* Predicting Protein Functions Based on Heterogeneous Graph Attention Technique. *IEEE J Biomed Health Inform* **28**, 2408–2415 (2024).
50. Schwikowski, B., Uetz, P. & Fields, S. A network of protein–protein interactions in yeast. *Nature Biotechnology* **18**, 1257–1261 (2000).
51. Guangyuan F., Guoxian Y., Jun W. & Maozu G. Protein Function Prediction Using Positive and

Negative Examples. *Journal of Computer Research and Development* **53**, 1753–1765 (2016).

52. Yu, G., Fu, G., Wang, J. & Guo, M. Predicting irrelevant functions of proteins based on dimensionality reduction. *Scientia Sinica Informationis* **47**, 1349–1368 (2017).
53. You, R., Yao, S., Mamitsuka, H. & Zhu, S. DeepGraphGO: Graph neural network for large-scale, multispecies protein function prediction. *Bioinformatics* **37**, I262–I271 (2021).
54. Wu, K. *et al.* PSPGO: Cross-Species Heterogeneous Network Propagation for Protein Function Prediction. *IEEE/ACM Trans Comput Biol Bioinform* **20**, 1713–1724 (2023).
55. Zhao, C., Liu, T. & Wang, Z. PANDA2: Protein function prediction using graph neural networks. *NAR Genom. Bioinform.* **4**, (2022).
56. Cao, Y. & Shen, Y. TALE: Transformer-based protein function Annotation with joint sequence-Label Embedding. *Bioinformatics* **37**, 2825–2833 (2021).
57. Pan, T. *et al.* PFresGO: an attention mechanism-based deep-learning approach for protein annotation by integrating gene ontology inter-relationships. *Bioinformatics* **39**, (2023).
58. Gligorijević, V. *et al.* Structure-based protein function prediction using graph convolutional networks. *Nat. Commun.* **12**, (2021).
59. Littmann, M., Heinzinger, M., Dallago, C., Olenyi, T. & Rost, B. Embeddings from deep learning transfer GO annotations beyond homology. *Sci. Rep.* **11**, 1160 (2021).
60. Schütze, K., Heinzinger, M., Steinegger, M. & Rost, B. Nearest neighbor search on embeddings rapidly identifies distant protein relations. *Front. Bioinform.* **2**, 1033775 (2022).
61. Kaminski, K., Ludwiczak, J., Pawlicki, K., Alva, V. & Dunin-Horkawicz, S. pLM-BLAST: distant homology detection based on direct comparison of sequence representations from protein language models. *Bioinformatics* **39**, (2023).
62. Pantolini, L. *et al.* Embedding-based alignment: combining protein language models with dynamic programming alignment to detect structural similarities in the twilight-zone. *Bioinformatics* **40**, (2024).
63. Domingo-Fernández, D., Hoyt, C. T., Bobis-Álvarez, C., Marín-Llaó, J. & Hofmann-Apitius, M. ComPath: an ecosystem for exploring, analyzing, and curating mappings across pathway databases. *NPJ Syst. Biol. Appl.* **5**, 3 (2019).
64. Haendel, M. A. *et al.* A census of disease ontologies. *Annu. Rev. Biomed. Data Sci.* **1**, 305–331 (2018).
65. Soman, K. *et al.* Biomedical knowledge graph-optimized prompt generation for large language models. *Bioinformatics* **40**, (2024).
66. Peng, B. *et al.* Graph retrieval-Augmented Generation: A Survey. *arXiv [cs.AI]* (2024) doi:10.48550/ARXIV.2408.08921.
67. Wang, H. *et al.* Can language models solve graph problems in natural language? *arXiv [cs.CL]* (2023) doi:10.48550/ARXIV.2305.10037.
68. UniProt Consortium. UniProt: the Universal Protein Knowledgebase in 2025. *Nucleic Acids Res* **53**, D609–D617 (2025).
69. Szklarczyk, D. *et al.* The STRING database in 2023: protein-protein association networks and functional enrichment analyses for any sequenced genome of interest. *Nucleic Acids Res* **51**, D638–D646 (2023).
70. Orchard, S. *et al.* The MIntAct project--IntAct as a common curation platform for 11 molecular interaction databases. *Nucleic Acids Res* **42**, D358–63 (2014).

71. Oughtred, R. *et al.* The BioGRID database: A comprehensive biomedical resource of curated protein, genetic, and chemical interactions. *Protein Sci.* **30**, 187–187 (2021).

72. Aleksander, S. A. *et al.* The Gene Ontology knowledgebase in 2023. *Genetics* **224**, (2023).

73. Bairoch, A. The ENZYME database in 2000. *Nucleic Acids Res.* **28**, 304–305 (2000).

74. Altenhoff, A. M. *et al.* OMA orthology in 2024: improved prokaryote coverage, ancestral and extant GO enrichment, a revamped synteny viewer and more in the OMA Ecosystem. *Nucleic Acids Res* **52**, D513–D521 (2024).

75. Blum, M. *et al.* InterPro: the protein sequence classification resource in 2025. *Nucleic Acids Res* **53**, D444–D456 (2025).

76. Knox, C. *et al.* DrugBank 6.0: the DrugBank Knowledgebase for 2024. *Nucleic Acids Res* **52**, D1265–D1275 (2024).

77. Mendez, D. *et al.* ChEMBL: towards direct deposition of bioassay data. *Nucleic Acids Res.* **47**, D930–D940 (2019).

78. Kuhn, M., von Mering, C., Campillos, M., Jensen, L. J. & Bork, P. STITCH: interaction networks of chemicals and proteins. *Nucleic Acids Research* **36**, D684 (2007).

79. Cannon, M. *et al.* DGIdb 5.0: rebuilding the drug-gene interaction database for precision medicine and drug discovery platforms. *Nucleic Acids Res* **52**, D1227–D1235 (2024).

80. Kuhn, M., Letunic, I., Jensen, L. J. & Bork, P. The SIDER database of drugs and side effects. *Nucleic acids research* **44**, (2016).

81. Tatonetti, N. P., Ye, P. P., Daneshjou, R. & Altman, R. B. Data-Driven Prediction of Drug Effects and Interactions. *Science Translational Medicine* **4**, 125ra31 (2012).

82. Cai, M.-C. *et al.* ADReCS: an ontology database for aiding standardization and hierarchical classification of adverse drug reaction terms. *Nucleic Acids Res* **43**, D907–13 (2015).

83. Vasilevsky, N. A. *et al.* Mondo: Unifying diseases for the world, by the world. (2022) doi:10.1101/2022.04.13.22273750.

84. Piñero, J. *et al.* The DisGeNET knowledge platform for disease genomics: 2019 update. *Nucleic Acids Res* **48**, D845–D855 (2020).

85. Grissa, D., Junge, A., Oprea, T. I. & Jensen, L. J. Diseases 2.0: a weekly updated database of disease–gene associations from text mining and data integration. *Database* **2022**, baac019 (2022).

86. Landrum, M. J. *et al.* ClinVar: public archive of relationships among sequence variation and human phenotype. *Nucleic Acids Res* **42**, D980–5 (2014).

87. Gargano, M. A. *et al.* The Human Phenotype Ontology in 2024: phenotypes around the world. *Nucleic acids research* **52**, (2024).

88. Buniello, A. *et al.* Open Targets Platform: facilitating therapeutic hypotheses building in drug discovery. *Nucleic Acids Res* **53**, D1467–D1475 (2025).

89. Kafkas, Ş. *et al.* PathoPhenoDB, linking human pathogens to their phenotypes in support of infectious disease research. *Sci Data* **6**, 79 (2019).

90. Kanehisa, M., Furumichi, M., Sato, Y., Matsuura, Y. & Ishiguro-Watanabe, M. KEGG: biological systems database as a model of the real world. *Nucleic acids research* **53**, (2025).

91. Garcia-Alonso, L., Holland, C. H., Ibrahim, M. M., Turei, D. & Saez-Rodriguez, J. Benchmark and integration of resources for the estimation of human transcription factor activities. *Genome Res* **29**, 1363–1375 (2019).

92. Müller-Dott, S. *et al.* Expanding the coverage of regulons from high-confidence prior knowledge for accurate estimation of transcription factor activities. *Nucleic Acids Res* **51**, 10934–10949 (2023).

93. Han, H. *et al.* TRRUST v2: an expanded reference database of human and mouse transcriptional regulatory interactions. *Nucleic Acids Res* **46**, D380–D386 (2018).

94. Sheils, T. K. *et al.* TCRD and Pharos 2021: mining the human proteome for disease biology. *Nucleic Acids Res* **49**, D1334–D1346 (2021).

95. Davis, A. P. *et al.* Comparative Toxicogenomics Database's 20th anniversary: update 2025. *Nucleic Acids Res* **53**, D1328–D1334 (2025).

96. Moreno, P. *et al.* Expression Atlas update: gene and protein expression in multiple species. *Nucleic Acids Research* **50**, D129–D140 (2022).

97. Lonsdale, J. *et al.* The Genotype-Tissue Expression (GTEx) project. *Nature Genetics* **45**, 580–585 (2013).

98. Elnaggar, A. *et al.* ProtTrans: Towards cracking the language of life's code through self-supervised deep learning and high performance computing. *arXiv [cs.LG]* (2020) doi:10.48550/ARXIV.2007.06225.

99. Lin, Z. *et al.* Evolutionary-scale prediction of atomic-level protein structure with a language model. *Science* **379**, 1123–1130 (2023).

100. Dalla-Torre, H. *et al.* The Nucleotide Transformer: Building and evaluating robust foundation models for human genomics. *bioRxiv* (2023) doi:10.1101/2023.01.11.523679.

101. Řeh uřek, R. & Sojka, P. *Software Framework for Topic Modelling with Large Corpora*. (ELRA, 2010).

102. Peng, C. *et al.* CADA: phenotype-driven gene prioritization based on a case-enriched knowledge graph. *NAR Genom. Bioinform.* **3**, (2021).

103. Melidis, D. P. & Nejdl, W. Capturing Protein Domain Structure and Function Using Self-Supervision on Domain Architectures. *Algorithms 2021, Vol. 14, Page 28* **14**, 28–28 (2021).

104. Ali, M., Hoyt, C. T., Domingo-Fernández, D., Lehmann, J. & Jabeen, H. BioKEEN: a library for learning and evaluating biological knowledge graph embeddings. *Bioinformatics* **35**, 3538–3540 (2019).

105. Schwaller, P. *et al.* Mapping the space of chemical reactions using attention-based neural networks. *Nature Machine Intelligence 2021 3:2* **3**, 144–152 (2021).

106. Edera, A. A., Milone, D. H. & Stegmayer, G. Anc2vec: embedding gene ontology terms by preserving ancestors relationships. *Brief. Bioinform.* **23**, (2022).
